# Abstract rule generalization for composing novel meaning recruits the frontoparietal control network

**DOI:** 10.1101/2024.10.30.621101

**Authors:** Xiaochen Y. Zheng, Mona M. Garvert, Hanneke E. M. den Ouden, Lisa I. Horstman, David Richter, Roshan Cools

**Author notes:** Correspondence to Xiaochen Y. Zheng, Donders Centre for Cognitive Neuroimaging, Kapittelweg 29, 6525 EN, Nijmegen, The Netherlands.

## Abstract

The ability to generalize previously learned knowledge to novel situations is crucial for adaptive behavior, representing a form of cognitive flexibility that is particularly relevant in language. Humans excel at combining linguistic building blocks to infer the meanings of novel compositional words, such as “un-reject-able-ish”. How do we accomplish this? While recent research on compositional generalization in relational memory, action planning and vision strongly implicates a medial prefrontal-hippocampal network (Baram et al., 2021; Barron et al., 2020; Schwartenbeck et al., 2023), it remains unclear whether the same network supports compositional inference in language. To this end, we trained participants on an artificial language in which the meanings of compositional words could be derived from known stems and unknown affixes, using abstract affixation rules (e.g., “good-kla” which means “bad”, where “-kla” reverses the meaning of the stem word “good”). According to these rules, word meaning depended on the sequential relation between the stem and the affix (i.e., pre- vs. post-stem). During fMRI, participants performed a semantic priming task, with novel compositional words as either sequential order congruent (e.g., “white-kla”) or incongruent primes (e.g., “kla-white”), and synonyms of the composed meaning as targets (“black”). Our results show that the compositional process engages a broad temporoparietal network, including the hippocampus, while the composed meanings are represented in left-lateralized language areas. Notably, newly composed meanings were decodable already at the time of the prime. Finally, we found that the composition process recruits abstract structure rule representations in a lateral frontoparietal network, rather than the predicted medial frontotemporal network, perhaps because abstract rules in language are more readily formatted as production rules rather than as relational map structures.

## Introduction

The ability to generalize previously acquired information to novel scenarios is essential for adaptive behavior in a changing world. While this hallmark of human cognition underpins learning and problem-solving across various cognitive domains (Behrens et al., 2018; Dehaene et al., 2022; Frankland & Greene, 2020; Gärdenfors, 2004; Schwartenbeck et al., 2023), this capacity is particularly clearly illustrated by language. When encountering the novel word “un-reject-able-ish” for the first time, we can swiftly infer its meaning by generalizing from the known elements and integrating them according to abstract relational structure rules, such as the sequential arrangement of word parts. We excel at combining linguistic building blocks such as morphemes and words to form larger structures like phrases and sentences, thereby flexibly conveying an infinite array of thoughts and ideas. The generation of linguistic meaning relies not only on the constituent parts, but more importantly, also on the abstract relational structure rules based on which they are combined (Fodor, 1975; Fodor & Pylyshyn, 1988; Frege, 1892; Martin, 2016; Partee, 2008). Consider the sentences “The cat chased the mouse” and “The mouse chased the cat”. Despite sharing identical linguistic building blocks, they convey distinct meanings. What neural mechanisms enable us to infer novel compositional meaning based on such rules? Does our brain represent abstract relational structure rules to facilitate meaning generalization, and if so, which circuits are recruited?

In cognitive neuroscience, extensive research has been dedicated to understanding how our brain organizes knowledge to guide flexible behavior. An influential line of inquiry has focused on how this organization is achieved through learning simplified and abstract representations of the world, formatted as cognitive maps (Constantinescu et al., 2016; Moser et al., 2008; O’Keefe & Nadel, 1978; Solomon et al., 2019; Tolman, 1948; Zheng, Hebart, et al., 2024). These relational knowledge structures allow us to infer novel associations that have not been directly experienced, and to generalize those abstract structures to novel situations (Bein & Niv, 2023; H. Eichenbaum & Cohen, 2014; Piaget, 1929; Preston & Eichenbaum, 2013). In a recent study, Schwartenbeck et al., (2023) investigated the neural representations and mechanisms that enable compositional generalization in the domain of vision. Participants solved compositional problems by inferring the relational positions of building blocks in a visual silhouette (e.g., a building block on top of vs. below another building block). Using fMRI, they found generalizable, relational configurations of visual building blocks to be represented in a medial prefrontal-hippocampal network. The same network has been shown to be recruited during various other forms of generalization, ranging from discovering a shortcut in spatial navigation (Epstein et al., 2017; Jacobs et al., 2013; Moser et al., 2008; O’Keefe & Nadel, 1978; Tolman, 1948), to “joining the dots” between events (Barron et al., 2013, 2020; Garvert et al., 2023; Morton et al., 2020), and to inferring unknown relationships in social contexts (Park et al., 2020, 2021). It has been recently proposed that neural cognitive map-like representations in circuitry connecting the hippocampus with the medial frontal cortex can serve as a universal knowledge code for generalization and novel inference across multiple cognitive domains (Behrens et al., 2018; Bellmund et al., 2018; Stachenfeld et al., 2017; Whittington et al., 2018).

In the language sciences, the investigation of compositional generalization has however primarily implicated neural networks other than this medial prefrontal-hippocampal network. Compositionality – the ability to combine lexical building blocks to create linguistic meaning (Fedorenko et al., 2016; Gwilliams, 2020; Hagoort, 2019a, 2019b; Hagoort & Indefrey, 2014; Martin, 2020; Pylkkänen, 2019; Zaccarella et al., 2017; Zaccarella & Friederici, 2015) – is thought to rely on left-lateralized, language-specific networks, particularly in regions such as the left inferior frontal gyrus (Bozic et al., 2007; Bozic & Marslen-Wilson, 2010; Hagoort, 2005, 2016; Leminen et al., 2019; Nevat et al., 2017) and the left anterior temporal lobe (Baron & Osherson, 2011; Brennan et al., 2012; Flick et al., 2018; Pylkkänen, 2019). This suggests that compositional inference in language might engage neural system distinct from those involved in compositional processes in relational memory, action planning and vision, challenging the notion that hippocampal-based representational codes are truly domain-general.

In the current pre-registered fMRI study, we aimed to test this hypothesis by investigating the neural mechanisms underlying the ability to infer novel compositional word meanings based on relational structure rules. Specifically, we aimed to assess whether relational structure-based composition in language recruits the medial prefrontal-hippocampal network that has also been implicated in action planning, visual composition and relational memory (Baram et al., 2021; Barron et al., 2020; Schwartenbeck et al., 2023). To this end, we employed a recently developed artificial language learning paradigm where participants generalize abstract relational structure rules to infer novel compositional meanings (Zheng, Petukhova, et al., 2024). In this task, participants infer abstract affixation rules from linguistic exemplars, then use these rules to derive the meanings of novel compositional words. According to these rules, word meaning depends on the sequential relation between the stem and the affix (i.e., pre- vs. post-stem). The paradigm was designed to capture (i) the observation that sequential order plays a key role in compositionality in natural language (Beyersmann & Grainger, 2023; Crepaldi et al., 2013, 2016), but furthermore, also (ii) the relational structure-dependent nature of compositional generalization in non-linguistic domains associated with hippocampal-medial frontal cortical circuitry (Baram et al., 2024; Barron et al., 2020; Garvert et al., 2023; Morton et al., 2020; Park et al., 2021; Schwartenbeck et al., 2023). Our results show that newly inferred meanings are represented in language-specific left frontal regions (Hagoort, 2005, 2016; Nevat et al., 2017). Intriguingly, this generalization process recruits abstract rule representations in a lateral frontoparietal network that also represents abstract task representations for generative action selection (Vaidya & Badre, 2022), rather than the predicted medial prefrontal-hippocampal network, often associated with relational structure coding.

## Results

### Generalization of Abstract Rules for Novel Meaning Inference

To quantify participants’ ability to construct compositional word meaning by generalizing abstract structure rules to unknown context, we employed an experimental paradigm where participants learned an artificial language featuring various compositional rules (Zheng, Petukhova, et al., 2024). During a pre-scanning training phase, 30 healthy participants were exposed to pairs of compositional pseudo-words along with their experimentally assigned meanings (Figure 1A). Each of these compositional pseudo-words comprises a known stem (e.g., “good” in “good-kla”) and an unknown affix (e.g., “kla”). We manipulated the mapping of the meaning to the affix based on their sequential position: e.g., “-kla” as a suffix means “the opposite”, whereas “kla-” as a prefix means “young version”. These position-dependent affixation rules allowed participants to compose unique meanings based on different sequential combinations of the affixes with the stems. Crucially, while the participants could infer the affixation rules from the exemplars, these rules were never made explicit to them. Participants learned the meanings of all pseudo-words in four training blocks, evidenced by ceiling level performance on a subsequent memory task where they were instructed to recall the meanings of these words (mean_accuracy_ = 98.2 %, SD = 3.1%, Figure 1B).

**Figure 1.**
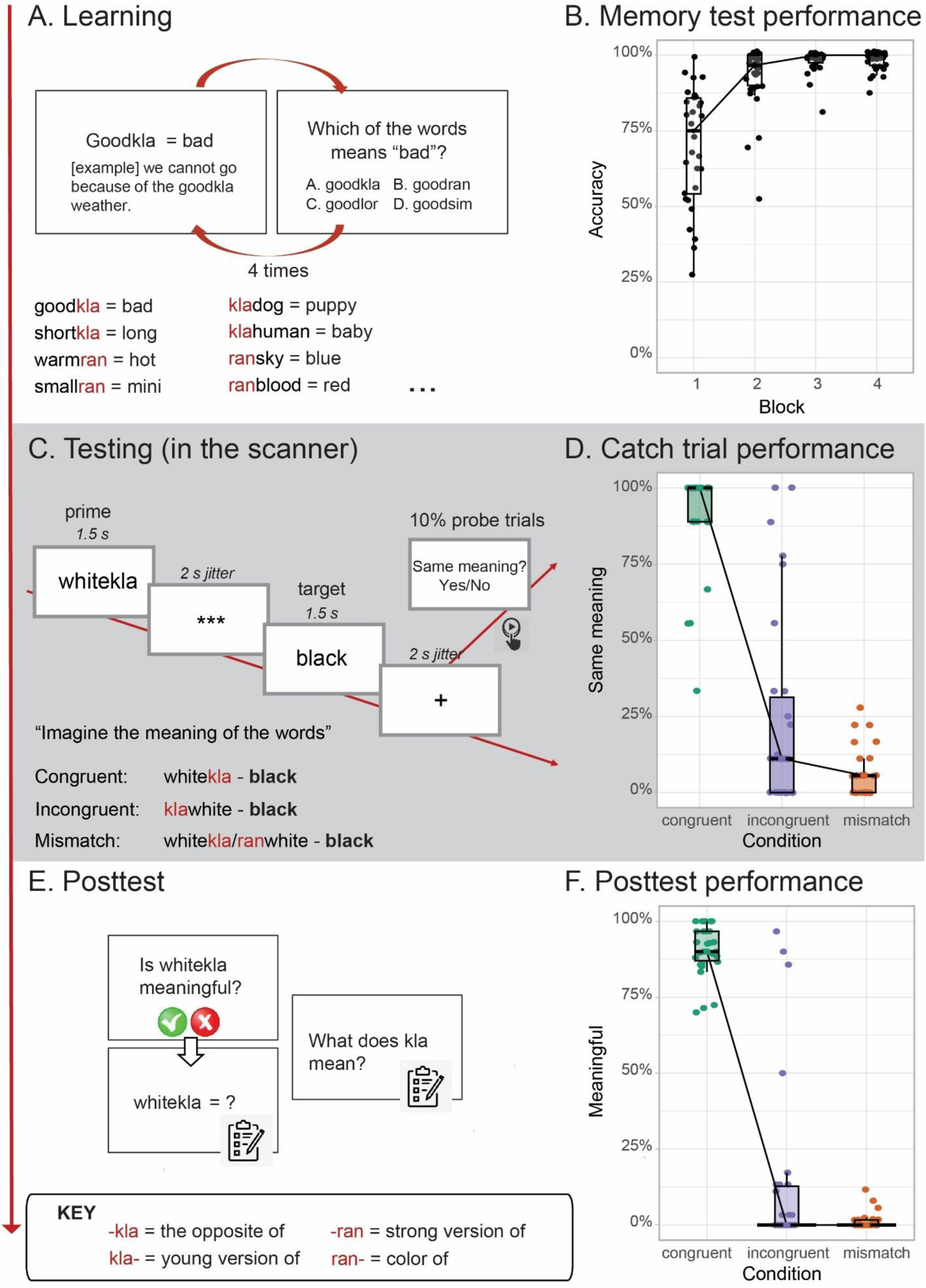
Experimental design (A, C, E) and behavioral results (B, D, F). (A) Participants learned and memorized artificial, compositional words. These compositional pseudo-words consisted of a known stem and an unknown affix. The affix alters the word meaning depending on its position (pre- vs. post-stem). Importantly, the relational structure rules were never made explicit to the participants. (B) Box plots of participant’s choice performance in a memory task, where they recall the meaning of the learned pseudo-words. (C) We tested participants’ knowledge with novel, compositional pseudo-words using an fMRI adaptation paradigm, in which the prime pseudo-words were always followed by a target, real word. The prime pseudo-words were either congruent or incongruent in terms of the relational structure rules, or totally mismatched in meaning regardless of order. The target word was always a matched synonym to the congruent prime word. (D) Boxplots of participants’ responses in the fMRI task across three experimental conditions, on the 10% probe trials on which they indicated with a left or right button press whether the prime words did or did not match the target words in terms of their meaning. (E) After the fMRI session, we explicitly asked participants to evaluate the meaningfulness of the novel compositional words. (F) Boxplots of participants’ responses in the posttest across three experimental conditions, where they indicated whether the pseudo-words are meaningful or not. The actual stimuli used in the experiment were in participants’ native language, Dutch (Supplementary Material 2). For (B, D, F), the thick horizontal line inside the box indicates the group median, and the bottom and top of the box indicate the group-level first and third quartiles of each condition. Each dot represents one participant. The black lines connect the group median across conditions.

To test participants’ knowledge of the abstract structure rules, we presented a new set of compositional pseudo-words that they had never encountered before (e.g. “white-kla” and “kla-white”) and asked them to imagine the meanings of the words, while recording fMRI. These novel words could be either congruent with the sequential order rule they had learned (e.g. for “white-kla”, “-kla” as suffix means “the opposite”, and the opposite of white is “black”), or incongruent (e.g. for “kla-white”, “kla-” as a prefix means “the young version of”, while it is much more difficult to infer the meaning of the young version of white). We presented the pseudo-words (“primes”) in pairs with their synonyms (“targets”), while the synonyms were always matched to the congruent condition (e.g., “black” was presented after “white-kla” and also after “kla-white”, Figure 1C). After 10% of the targets, participants were presented a probe question, asking whether the meaning of the target word was the same as that of the preceding, pseudo-word prime. Analysis of participants’ responses to these probe trials showed significantly higher probability of meaning-match responses in congruent (mean = 90.7%, SD = 16.5%) than incongruent trials (mean = 23.8%, SD = 32.3%; β = 4.52, SE = 0.78, z = 5.80, p < .001; Figure 1D), evidencing their reliance on the relational structure rules for inference. These results were validated in a posttest administered outside the scanner, where participants explicitly indicated whether they considered the novel pseudo-words that they had seen during the preceding MRI scan to be meaningful or not (Figure 1E, 1F, Supplementary Materials 1). We further included mismatched pseudo-words as a control condition, where the stems were combined with alternative affixes, so that the meaning of the compositional pseudo-words did not match the target synonyms regardless of the position of the affix (e.g., ran-white = the color of white ≠ black; white-ran = the extreme version of white ≠ black). Participants did not consider these pseudo-words to match the meaning of the synonym, and their responses in this mismatch condition did not differ from those in the incongruent condition (Supplementary Materials 1). For clarity, we will now focus on the contrast between congruent versus incongruent conditions.

Together, these behavioral results demonstrate that participants were able to efficiently compute novel compositional meaning by generalizing previously learned structure rules to novel situations. The setup was optimized for capturing neural adaptations in fMRI, and allowed us to assess activity in neural circuits commonly associated with novel inference and abstract rule-based generalization.

### Compositional Meaning Representations in Language-Specific Frontal Regions

To explore the neural mechanisms associated with the compositional process, we compared fMRI BOLD responses during congruent versus incongruent primes, when participants first encountered the novel pseudo-words. The results showed greater activity for incongruent than congruent primes in multiple temporal and parietal areas, including the precuneus, the postcentral gyrus, and the lingual gyrus (Figure 2A, Supplementary Table S3-1). To uncover the neural representations of the composed meanings, we exploited the phenomenon of fMRI adaptation (also termed as “repetition suppression”; Barron et al., 2016; Grill-spector et al., 2006). Because the repeated activation of the same population of neurons leads to a suppressed response, we reasoned that in areas representing word meanings the response of the population of neurons should be suppressed upon repeated exposure to the same word meaning. In our task, neural signals at the time of the target would be suppressed to a greater degree when that target was preceded by a congruent prime word that shared the same meaning, compared with an incongruent prime. Analysis of the contrast of interest (incongruent – congruent targets) revealed greater adaptation of fMRI activity in a broad network of brain regions, including the (pre)cuneus, the postcentral gyrus, the middle frontal gyrus (Figure 2B; Supplementary Table S3-2), as well as the left inferior frontal cortex (Figure 2D; p_FWE_ < .001, K_E_ = 1118, Z_max_ = 4.47, MNI coordinates of the peak = [−50, 33, 10]), a region often associated with deriving new and complex meaning from the lexical building blocks (Hagoort, 2005, 2016; Nevat et al., 2017; Weber et al., 2016; W. Zhang et al., 2022). Notably, the reverse contrast revealed greater activation during the congruent versus incongruent condition in the striatum, both at the time of the prime and the target (Figure 2C, 2D; Supplementary Materials 3A).

**Figure 2.**
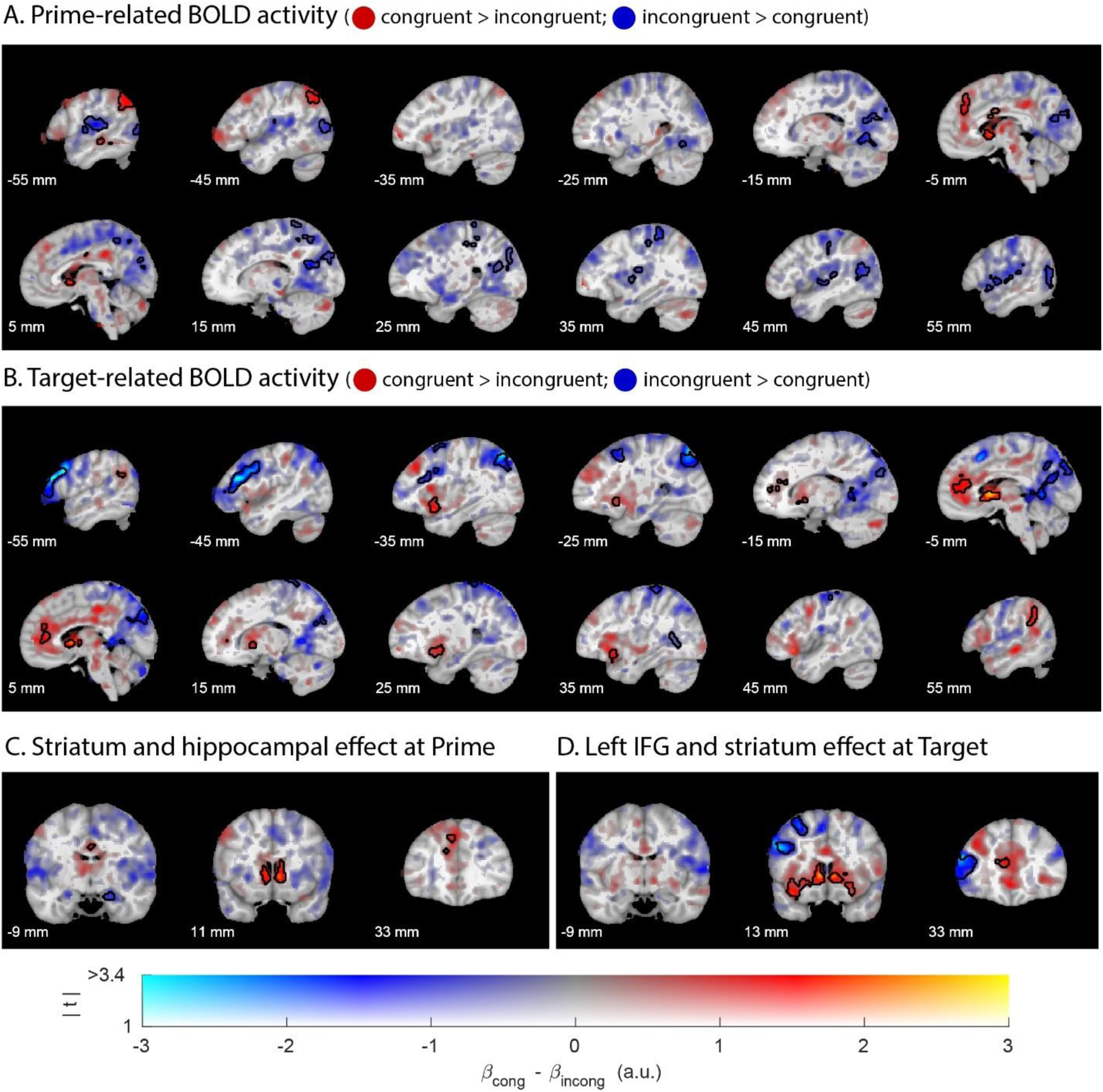
Univariate fMRI effects of novel meaning composition (prime-related activity) and its representational outcome (target-related activity). In red: congruent > incongruent; in blue: incongruent > congruent. (A) fMRI effects of congruent versus incongruent prime-related BOLD activity engages a broad temporoparietal network. (B) fMRI effects of congruent versus incongruent target-related BOLD activity (in blue: fMRI adaptation) reveal composed meaning representations in the left inferior cortex. (C) Prime-related effects of interest in the hippocampus and the striatum. (D) Target-related fMRI adaptation effects in the left IFG (in blue), but absent in the hippocampus. The hue indexes the sign and size of the contrast parameter estimate (congruent minus incongruent), and the opacity indexes the magnitude of the associated t values. Significant clusters (cluster-level corrected, FWE, p < .05) are encircled in solid contours. All coordinates are provided in the MNI space.

Based on previous work on nonlinguistic composition and generalization, particularly in the domain of relational memory (Barron et al., 2020; Garvert et al., 2023; Jacobs et al., 2013; Morton et al., 2020; Park et al., 2021; Schwartenbeck et al., 2023), we hypothesized that the process of composing novel meanings elicits activity in a circuit connecting the hippocampal formation with the medial prefrontal cortex. To test the engagement of this network, we performed additional analyses within small volumes of interest in the medial prefrontal cortex (mPFC; defined functionally based on Schwartenbeck et al., 2023) and the hippocampal formation (defined anatomically, Supplementary Materials 5). This analysis revealed greater activity during incongruent than congruent primes in the hippocampal formation (Figure 2C, p_FWE_ = .045, K_E_ = 26, Z_max_ = 3.72, [17, −9, −20]), perhaps reflecting greater effort to resolve the generalization-based composition challenge during incongruent than congruent primes. There was no evidence for effects of prime type in the mPFC (no suprathreshold clusters found after small volume correction, SVC). In contrast to our hypothesis, during target words there was no evidence for differences in fMRI adaptation in the hippocampal formation after congruent versus incongruent primes (Figure 2D, p_FWE_ = .203, K_E_ = 6, Z_max_ = 3.50, [29, −38, 4], SVC). Moreover, the activity in mPFC was actually greater during targets following congruent than incongruent primes (Figure 2B; p_FWE_ = < .001, K_E_ = 628, Z_max_ = 4.26, [−12, 41, 20]). Thus, these results do not substantiate our hypothesis that novel word meanings are represented in the medial prefrontal-hippocampal network.

To probe the role of the language network in meaning inference, we performed additional SVC for the left anterior temporal lobe and the left inferior frontal gyrus (left IFG). Apart from the above-mentioned fMRI adaptation effect in left IFG for incongruent versus congruent targets, we found no further evidence for these regions in meaning composition (Supplementary Material 3B). Moreover, a supplementary analysis of contrasts between the congruent and mismatch conditions revealed qualitatively similar patterns of effects, thus substantiating these univariate analyses that focused on the congruent vs incongruent contrast (Supplementary Material 3C).

In sum, the process of composing meaning based on abstract structure rules, as measured in terms of prime-related BOLD signal, was associated with neural activity in a broad temporoparietal network, including the hippocampal formation. In addition, the representational outcome of this compositional process surfaced as fMRI adaptation of target-related neural activity in areas often associated with language processing, including the left inferior frontal cortex. Finally, successful meaning composition at target was accompanied by BOLD change in the striatum and the mPFC, perhaps reflecting intrinsic reward signaling.

### Abstract Relational Rule Representations in Lateral Frontoparietal Network

The fMRI adaptation effect at the time of the congruent vs incongruent *target* likely occurred because participants already composed the newly inferred word meaning using the relational structure rules when they first encountered the primes. We tested this hypothesis directly by investigating the representations of both these relational structure rules and the newly inferred word meanings at the time of prime, using multivariate representational similarity analyses (RSA; Kriegeskorte et al., 2008; Nili et al., 2014). Consider the compositional pseudo-word “white-kla”: To compose its meaning, participants would represent the *rule* (“-kla” means “the opposite of”); Moreover, provided successful composition, they would also represent the composed *meaning* (“white-kla” means “black”). RSA allowed us to capture the relevant neural representations by computing the neural representational dissimilarity matrices (neural RDMs) based on prime-related fMRI activity for each pseudo-word, analyzed through a whole-brain searchlight. Next, we assessed whether these neural RDMs were explained by model RDMs that capture the similarities between these pseudo-words as a function of either their composed meaning or the rule that was used to compose them. Before exploring the two types of representations of interest, we first validated our RSA approach by constructing an RDM based on a model of the visual similarity of the target words to predict target-related neural activity. As anticipated, this RSA revealed significant shared representational geometry in the visual cortex, indicating that visual aspects of the word forms are represented in visual area (Supplementary Material 4A).

We then constructed a meaning model (Figure 3A) to capture the representations of the newly composed word meanings, in which word meanings are arranged by their semantic similarities (e.g., “black”, derived from “whitekla”, is more similar to “dark” - derived from “lightkla” - compared with “happy” - derived from “sadkla”; based on word embeddings, Mandera et al., 2017). We reasoned that the similarity of BOLD patterns in brain areas that represent the newly constructed word meanings (e.g., white-kla” means black) should reflect the semantic similarity of the composed words (e.g., “black”), as derived from this word meaning model. RSA using a whole-brain searchlight approach, showed that the newly constructed meanings were represented in left lateralized language-related areas (Figure 3C), including the left inferior frontal cortex (p_FWE_ = .001, K_E_ = 98, Z_max_ = 4.06, [−48, 17, 16]) and angular gyrus (p_FWE_ < .001, K_E_ = 377, Z_max_ = 4.09, [−42, −52, 48], Supplementary Table S2-2). The pattern of these RSA effects, computed from prime-related activity, overlapped greatly with the pattern of RSA effects, computed from target-related neural activity (predicted by the same target-meaning model; Supplementary Material 4B). It is also worth noting that this left inferior frontal area overlapped greatly with the left frontal cluster yielded by the univariate fMRI adaptation analysis, that is the comparison between congruent vs incongruent targets. Importantly, this prime-related meaning representation was not captured by an alternative meaning model which described the similarities between stem meanings (e.g., “white” in “whitekla” is more similar to “light” in “lightkla”, compared with “sad” in “sadkla”; Kendall’s τ_stem-target_ = .13; all cluster-level ps > .841; Supplementary Material 4C). Together, this RSA analysis demonstrates that the representation of the novel meaning was already decodable at the time of pseudo-word primes.

**Figure 3.**
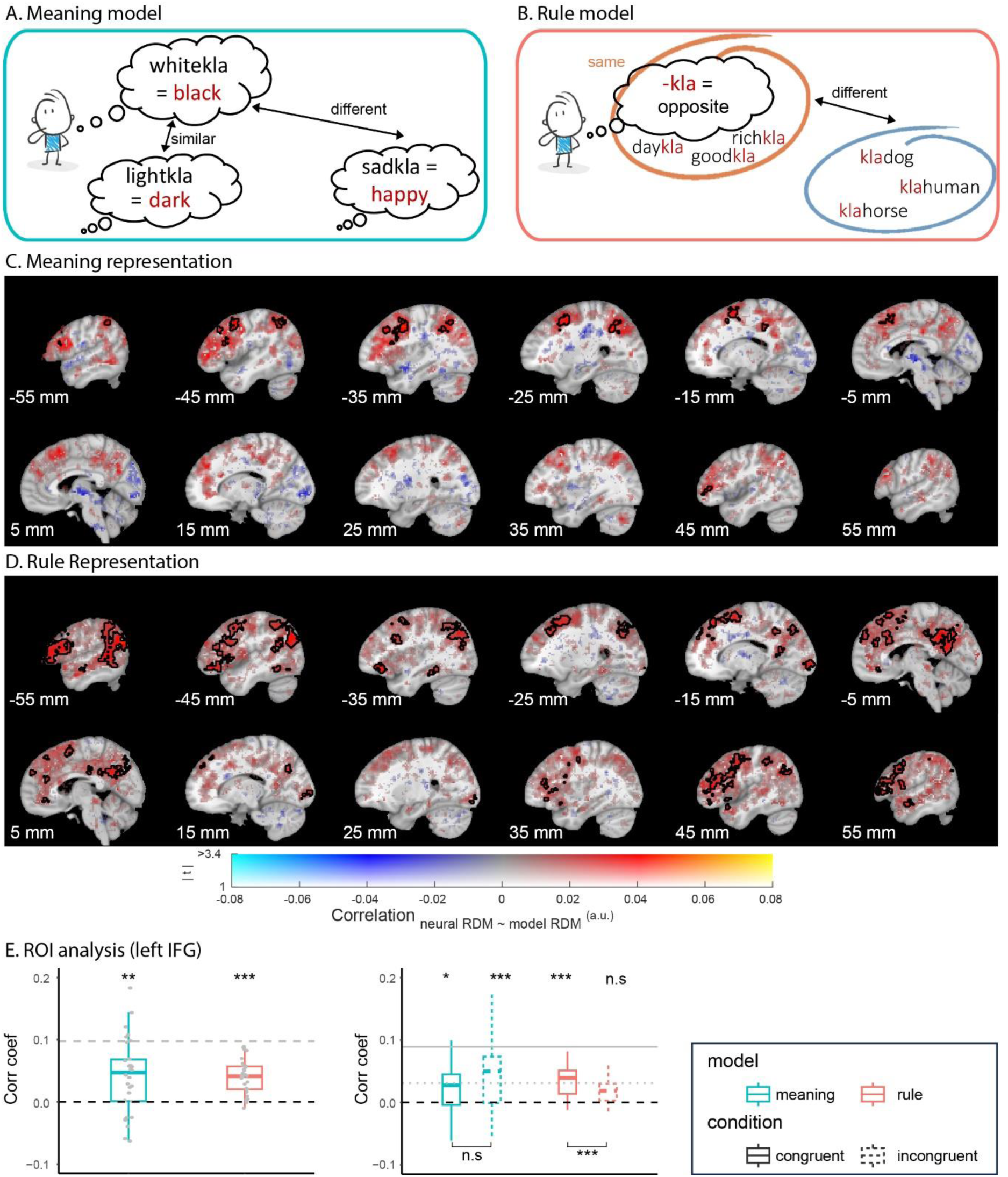
Representational similarity analysis of meaning and rule representations. (A) A distance-based meaning model in which word meanings are arranged by their similarities. (B) A binary-coded rule model in which all the compositional pseudo-words ending with the same affix (e.g., “-kla”) are more similar to each other, compared with words with a different affix (e.g., “kla-”, “-ran”, or “ran-”). (C, D) Whole-brain searchlight RSA outcome using the meaning model and the rule model, respectively. Effects are shown from an analysis in which congruent and incongruent conditions were combined. The hue indexes the sign and size of the correlation coefficient, and the opacity indexes the magnitude of the associated t values. Significant clusters (cluster-level corrected, FWE, p < .05) are encircled in solid contours. All coordinates are provided in the MNI space. (E) ROI-based RSA of meaning (green) and rule (red) representations, extracted from the left inferior frontal gyrus, for congruent and incongruent conditions combined (left panel) and separately (right panel). The gray horizontal line indicates the noise ceiling of the data extracted from the respective ROIs, computed using a leave-one-out approach. The noise ceiling estimates the maximum performance that can be achieved on the measure under ideal conditions, assuming no noise or variability. Asterisks indicate the statistical significance based on partial correlations: ***: < .001; **: < .01; *: < .05; n.s: not significant.

Next, we constructed a rule model (Figure 3B) to capture the representation of the relational rules, in which all compositional pseudo-words ending with “-kla” are more similar to each other than the pseudo-words with a different affix (e.g., “kla-”, “-ran”, or “ran-”). We reasoned that for brain regions where the abstract relational structure rules are represented, their BOLD pattern should be best explained by the rule model (a 30*30 binary-coded distance matrix where the rules are either the same, or different). Whole-brain searchlight RSA revealed representations of the newly derived abstract rules in a bilateral frontoparietal network (Figure 3D), including dorsolateral prefrontal cortex (p_FWE_ < .001, K_E_ = 1639, Z_max_ = 4.59, [59, 7, 14]), middle temporal gyrus (p_FWE_ < .001, K_E_ = 4619, Z_max_ = 5.48, [−54, −52, 0]), but also medial prefrontal cortex (p_FWE_ < .001, K_E_ = 273, Z_max_ = 5.01, [−6, 25, 38], Supplementary Table S2-2). These areas are commonly implicated in the representation of task-state spaces and abstract rules in working memory for goal-directed action planning (Cole, Reynolds, et al., 2013; Cole & Schneider, 2007; Dosenbach et al., 2007; Harding et al., 2015; Nee, 2021; Spreng et al., 2010; Vaidya & Badre, 2022; Zanto & Gazzaley, 2013).

In contrast to our hypothesis, there was no evidence for either meaning or rule representations in the hippocampal formation, also not when reducing the search volume to an anatomically defined ROI (Supplementary Material 4D). Interestingly, ROI-based RSA in the left IFG revealed that the rule representation was stronger for the congruent than the incongruent conditions (Figure 3E, Mean_cong_ = .03, SD = .03; Mean_incong_ = .006, SD = .03; Paired t29 = 3.51, p = .001). There was no significant effect, however, of congruency condition on the meaning representation (Mean_cong_ = .02, SD = .05; Mean_incong_ = .04, SD = .05; Paired t29 = −1.82, p = .080; Supplementary Material 4D, 4E).

In sum, our results show that the online generalization of abstract rules for composing novel word meaning engages a broad temporoparietal network including the hippocampus, with the newly composed words being represented in language-specific regions. During compositional generalization, both the newly inferred word meanings and the abstract rules can be decoded from a lateral frontoparietal control network.

## Discussion

Our brain learns and abstracts generalizable knowledge to make inference and adapt to novel situations (Al Roumi et al., 2019; Behrens et al., 2018; Dehaene et al., 2022; Frankland & Greene, 2020; Gärdenfors, 2004; Liu et al., 2019; Sablé-Meyer et al., 2022; Schwartenbeck et al., 2023). Here, we investigated this flexibility of human cognition in composing novel word meaning based on previously learned abstract rules. Using fMRI adaptation, we demonstrated that newly inferred meanings are represented in the left inferior frontal cortex (IFC), a key region for constructing linguistic meaning (Nevat et al., 2017; Weber et al., 2016; W. Zhang et al., 2022). While we interpret the reduced neural activity for congruent versus incongruent targets as neural adaptation, it is also possible that this difference reflects prediction error in response to incongruent targets (see also Weber et al., 2016). Although our current paradigm cannot distinguish prediction error from neural adaptation (cf. Todorovic & de Lange, 2012), both interpretations support the conclusion that novel word meanings must have been composed “on the fly” and be represented at the time of the target. Furthermore, the RSA-derived finding that novel meaning representations were decodable from the IFC reinforces the conclusion that the IFC represents the composed meanings themselves rather than representing prediction error. It is remarkable that these newly constructed meanings were already represented at the time of the prime, and demonstrates that the current procedure provides a novel approach for quantifying the strength of the covert products of novel compositional inference.

While we observed hippocampal engagement during the composition *–* specifically when participants encountered incongruent words compared with congruent ones, potentially reflecting greater efforts to resolve generalization-based composition challenges – there was no clear evidence that the newly composed words themselves were represented in the hippocampus or the mPFC. This contrasts with our original prediction, given the critical role of medial prefrontal-hippocampal network in relational structure-based inference and generalization across domains such as vision, memory, and planning (Baram et al., 2021; Barron et al., 2013, 2020; Behrens et al., 2018; Bellmund et al., 2018; Garvert et al., 2023). One possible explanation lies in the representational nature of the abstract affixation rules used in our task: The medial prefrontal-hippocampal network is thought to maintain task knowledge in a flexible cognitive map, with these map-like representations supporting novel shortcuts or connections that are never experienced (Barron et al., 2020; Garvert et al., 2023; Jacobs et al., 2013; Morton et al., 2020; Park et al., 2021; Schuck et al., 2016; Schuck & Niv, 2019; Schwartenbeck et al., 2023). In hindsight, the abstract rules in our task seem unlikely to be formatted in terms of such a relational map; instead, they are more likely to be formatted as production rules (e.g., “if ‘kla’ is affixed at the end of a word, then it reverses the meaning of the word”).

To assess the neural locus of these abstract relational rule representations, we took an RSA approach. Results revealed rule representations in a lateral frontoparietal network, including the dorsolateral prefrontal cortex (DLPFC) and lateral parietal cortex. These regions have previously been associated with the learning and usage of abstract task representations during cognitive control (Badre et al., 2010; Cole, Laurent, et al., 2013; A. Eichenbaum et al., 2020; Ito et al., 2022; Loose et al., 2017; Nee, 2021; Reverberi et al., 2012; Tomov et al., 2018; Woolgar et al., 2011). This finding concurs generally with recent findings from an imaging study in which participants generalized (value-based) information across contexts based on learned abstract relationships that could also be conceptualized as being formatted as production rules (“if in context A, then category A+ gives more rewards”, Vaidya et al., 2021). These abstract production rules were also found to be represented in a similar frontoparietal control network, with flexible switching between rules facilitating efficient cognitive control.

Notably, the brain region identified as representing newly composed meanings, the left IFC, is anatomically close to the DLPFC, a core area of the aforementioned cognitive control network associated with structure rule abstraction and novel inference (Badre et al., 2010; Bartolo & Averbeck, 2021; Braver et al., 2002; Monti et al., 2007; Morton et al., 2020; Ramawat et al., 2022; Wallis et al., 2001). This might suggest a shared system for linguistic and nonlinguistic generalization, with the left hemisphere playing a more prominent role in facilitating communication within the language network (Fiebach et al., 2005; Hagoort, 2016). This interpretation is further supported by our findings of left-lateralized parietal involvement in abstract rule representations, such as the angular gyrus (see also W. Zhang et al., 2022).

When comparing congruent versus incongruent meaning compositions, we observed increased neural activity in the striatum, a region often implicated in reward processing (Knutson et al., 2000, 2001). This finding is surprising because we did not provide participants with reward feedback regarding the accuracy of their semantic judgments; they were not informed which words were congruent and therefore meaningful. While we must be cautious of reverse inference, this finding suggests that the process of generating meaning itself may provide an intrinsic reward. This aligns with the idea that internal rewards can facilitate the learning of grammar and new word meanings (Bains et al., 2024; Nevat et al., 2017; Ripollés et al., 2014; Ullman, 2016).

In sum, we introduced an artificial language learning paradigm for investigating cognitive flexibility and novel inference involved in linguistic composition. We leveraged the fact that participants can infer novel, compositional meanings on the fly, based on previously learned structure rules, and demonstrated that we can capture this ability using our procedure, as we have shown previously (Zheng, Petukhova, et al., 2024). Using fMRI, we demonstrated that abstract rule generalization for composing novel meaning recruits the frontoparietal control network and that the covert mental products of the compositional process can be decoded from the frontal cortex at the time of composition. The obvious next question is whether this paradigm can be leveraged to unravel the temporal dynamics of meaning composition. Future studies could use MEG to explore, for example: (1) whether word meaning representation in the left IFC reflects the transition from the stem word to the composed target word; (2) whether the representation of the presented structure rule shifts to its reversed-order rule when the latter results in successful composition.

## Methods

The study was approved by the local ethics committee (METC Oost-Nederland, 2014/288) and conducted in accordance with the Declaration of Helsinki. The study was pre-registered at AsPredicted (https://aspredicted.org/mk5i2.pdf).

### Participants

Given missing priors for this novel task, sample size was estimated based on 80% power to detect a median-size effect (Cohen’s d = 0.5) at an alpha of 0.05. We collected data from 43 right-handed, healthy Dutch native speakers (Mean_age_ = 23.1, SD_age_ = 4.3, range 18-33, 27 women, 15 men, 1 other). All participants had normal or corrected-to-normal vision. No participants reported to have any current or previous psychiatric or neurological disorders, nor MRI contraindications, such as unremovable metal parts in the body and claustrophobia. All participants provided written consent and received monetary compensation. Seven participants were excluded due to various reasons, including scanner failure (N = 1), poor fMRI data quality (N = 3, see criteria in *MRI data acquisition and preprocessing*), falling asleep in the scanner (N = 1), or fail to learn to generalize the structure rules (N = 4; 2 of which overlap with the ones with poor fMRI data, see criteria in *Behavioral analysis*). In addition, 6 participants were excluded due to an error in the stimuli list in the scanning session. This left us with a final sample of 30 participants (Mean_age_ = 23.0, SD_age_ = 3.5, range = 18-30, 19 women, 11 men).

### Experimental Paradigm

#### Materials

For the pre-scanning training phase, we constructed 30 compositional pseudo-words consisting of a known stem and an unknown affix. While we illustrate the design using English examples, the actual materials were in participants’ native language, Dutch. The affix altered the word meaning depending on its position and led to unique compositional meanings in different sequential combinations with the stems. We made sure that this sequential order rule could not be explained by the word class of the stems by misaligning the transitions (e.g., “-kla” turns an adjective to an adjective, whereas “kla-” turns a noun into a noun, however the same rule does not hold for affix “ran”, where both affix positions turn a noun into an adjective). In total, six affix meanings were selected, each with five exemplar words. These six affix meanings were assigned to three affix forms as prefix and suffix, respectively. The meaning-form assignment was counterbalanced across participants. In this way, we ensured the fMRI effects were not confounded by visual input. We also made sure that the compositional words do not resemble existing Dutch words.

For the testing phase in the scanner, a separate set of compositional pseudo-words (i.e., not presented in the learning) was created using another 30 stems. These pseudo-words were manipulated using the same affixation rules. Specifically, we manipulated the way in which the affixes are attached to the new stems, with each stem appearing in three different experimental conditions: (1) order-congruent: affixes are attached at the end, or at the beginning of the stem so that the novel compositional word is meaningful, given the position of the affix relative to the stem (e.g., white-kla = the opposite of white = black); (2) order-incongruent: affixes are attached at the end or beginning of the stem so that the novel compositional word is not meaningful, if participants had extracted the position-dependent rules from training (e.g., kla-white = the small version of white = meaningless). We made sure that, in this case, there was no possibility of inferring any meaning for the incongruent words using the sequential order rule; (3) mismatch: the stems are combined with alternative affixes, so that the novel compositional words are meaningless, regardless of the position of the affix (e.g., ran-white = “the color of” white = meaningless; white-ran = “the extreme version of” white = meaningless). Note that the items in the mismatch condition were twice as many as the congruent or the incongruent condition, due to the counterbalancing of the affix positions (e.g., white-kla is a congruent word, kla-white is an incongruent word, but both ran-white and white-ran are mismatch words). For all pseudo-words in the training phase and all order-congruent words in the testing phase, we made sure that there is an existing Dutch synonym which is not a complex word itself, to avoid a direct mapping of affix meanings. This synonym was paired with pseudo-words in all three conditions as the target word, during the scanning session. Given the uniquely powerful compositionality of language, it turned out to be impossible to ensure that all mismatch pseudo-words were meaningless. Instead, we ensured that in those rare cases, the meaning did not correspond to the target word. See Supplementary Material 1 for a full set of stimuli used for training and testing.

#### Procedure

Both the pre-scanning training and posttest were carried out in a sound-proof booth, whereas the scanning session was carried out in the MRI room. The experiment was run using the software Presentation (Version 20.2, Neurobehavioural System Inc, Berkeley, U.S.). A schematic diagram of the experiment is provided in Figure 1.

##### Pre-scanning training

Participants studied the training set of 30 compositional pseudo-words in a self-paced manner. Every compositional word was presented together with its synonym meaning and an example sentence using the word in context, till a maximum of 15 s or participants pressing to continue. After viewing all the words, participants completed a multiple-choice test where on each trial, they were given a synonym meaning and asked to choose a matched compositional word (e.g., “which of the words means ‘bad’? A. goodkla; B. goodran; C. goodlor; D. goodsen”). Each compositional word was presented once in a learning block and once in a memory test. All the words were presented in a pseudorandom order, with the same affix form or affix position repeated on no more than three consecutive trials. The learning blocks and memory tests were interleaved and repeated for 4 times, with 30 trials per block.

##### Scanning session

Next, participants went through a testing session in the MRI scanner. They were presented with the testing set of compositional pseudo-words (“prime”), paired with their matched or unmatched target synonyms (“target”). Participants were asked to imagine the meaning of the words presented on the screen.

Each prime word was presented on the screen for 1500 ms, followed by a jittered screen of “***”. The target word was then presented on the screen for 1500 ms, followed by another jittered screen of a fixation. Then the next trial started. Both jittered intervals were generated from a truncated exponential distribution with a mean of 2 s (range = 1.5 - 5 s). All prime words and target words were presented in black, in the center of a white screen. Each block started with a 2s fixation. All the pairs of prime and target were presented in a pseudorandom order, with the following requirements: (1) the same affix form or affix position repeated on maximal three consecutive trials; (2) the same condition repeated maximally for three consecutive trials; (3) The same stem was repeated at least five trials apart.

To ensure that participants paid attention to the prime words, we included a probe question on 10% of the trials where participants needed to indicate if the prime pseudo-word shares the same meaning as the target synonym word (“probe trials”). They responded by pressing the left (“yes”) or the right button (“no”) on the button box using their right index finger or middle finger, respectively. The probe question stayed on the screen for a maximum of 10s or until participants responded. The task then proceeded with a jittered fixation followed by the next trial.

The task consisted of three blocks in total, with each prime-target pair in each condition presented once in each block. Prior to going into the scanner, participants went through a practice block in the behavioral booth, where they got familiarized with the task and received feedback on their performance.

##### Posttest

In a posttest, participants were asked whether all the compositional pseudo-words they had seen in the scanning session were meaningful, and if so, what they meant. We also included the 30 pseudo-words presented in the learning set to assess participants’ memory performance. All the pseudo-words were presented in a pseudo-randomized order, using the same criteria as in the scanning session. At the end of the test, participants were asked explicitly to write down the meaning of the three sets of affixes.

At the end of the session, we also tested participants’ working memory capacity using an operation span task (Turner & Engle, 1989; Unsworth & Engle, 2005), where participants remembered letters while performing a mathematically calculation task (Mean = 49.3, SD = 12.7, range = 14-75). Additionally, we also collected information such as participants’ spoken foreign languages (Mean = 1.8, SD = 0.9, range = 1-4). This information were collected to explore the possible relationship between individual differences and learning of compositional rules (e.g., different strategies used, see Zheng, Petukhova, et al., 2024). However, we decided to skip the analysis given that participants did not show as much variance in their strategies as in the previous behavioral study.

The whole session took about 3 hours.

### MRI Data Acquisition and Preprocessing

#### Data Acquisition

The MRI experiment was performed on the institute 3T MAGNETOM Prisma[Fit] MR scanner (Siemens AG, Healthcare Sector, Erlangen, Germany) using a product 32-channel head coil. Out of the 30 participants in the final sample, 15 were scanned in a Prisma scanner and 15 were in a PrismaFit scanner. The assignments of participants were randomized. Despite the fact that the two scanners are theoretically the same, we additionally validated our results by including scanner as a second-level covariate. Our results hold still after excluding this potential covariance.

T2*-weighted blood-oxygen-level-dependent (BOLD) images were acquired in three blocks, recorded using a whole-brain multiband accelerated echo-planar imaging (EPI) sequence [TR, 1500 ms; TE, 39.6 ms; multiband acceleration factor, 4; flip angle, 75°; slice matrix size, 104 × 104; voxel size, 2.0 mm isotropic; FoV, 210 × 210 × 136 mm; bandwidth: 2090 Hz/px; echo spacing: 68 ms]. A high-resolution structural image (1 mm isotropic) was acquired using a T1-weighted 3D magnetization-prepared rapid gradient-echo sequence (MP-RAGE; TR, 2300 ms; TE, 3.03 ms; flip angle, 8°; FoV, 256 × 256 × 192 mm).

In addition, we recorded participants’ finger pulse using a pulse oximeter affixed to the second digit of the left hand. Respiration was measured using a respiration belt placed around the participant’s abdomen. These physiological data are shared in the associated dataset.

#### MRI Quality Control

The MRI quality control was performed using MRIQC 22.0.6. (Esteban et al., 2017). Means of framewise displacement (both in mm and in percentage of timepoints), temporal SNR, and DVAR for functional images were computed per participant per block based on the image quality metrics. Blocks with any of these values larger than 2.5 SD from the group mean were excluded (or smaller than 2.5 SD for temporal SNR). Individuals with two or more blocks excluded were also excluded from the dataset.

#### Preprocessing

All MRI data were preprocessed using fMRIPrep 21.0.2 (Esteban, Blair, et al., 2018; Esteban, Markiewicz, et al., 2018; RRID:SCR_016216), which is based on Nipype 1.6.1 (Gorgolewski et al., 2011, 2018; RRID:SCR_002502). Information in the two sections below (“Anatomical and Functional Data Preprocessing”) was retrieved directly from fMRIPrep.

In addition, we used Statistical Parametric Mapping 12 (SPM12; Wellcome Trust Centre for Neuroimaging, https://www.fil.ion.ucl.ac.uk/spm/) to spatially smooth the final preprocessed BOLD time series with a 6 mm FWHM kernel.

##### Anatomical Data Preprocessing

The T1-weighted (T1w) image was corrected for intensity non-uniformity (INU) with N4BiasFieldCorrection (Tustison et al., 2010), distributed with ANTs 2.3.3 (Avants et al., 2008, RRID:SCR_004757), and used as T1w-reference throughout the workflow. The T1w-reference was then skull-stripped with a Nipype implementation of the antsBrainExtraction.sh workflow (from ANTs), using OASIS30ANTs as target template. Brain tissue segmentation of cerebrospinal fluid (CSF), white-matter (WM) and gray-matter (GM) was performed on the brain-extracted T1w using fast (FSL 6.0.5.1:57b01774, RRID:SCR_002823, Zhang et al., 2001). Brain surfaces were reconstructed using recon-all (FreeSurfer 6.0.1, RRID:SCR_001847, Dale et al., 1999), and the brain mask estimated previously was refined with a custom variation of the method to reconcile ANTs-derived and FreeSurfer-derived segmentations of the cortical gray-matter of Mindboggle (RRID:SCR_002438, Klein et al., 2017). Volume-based spatial normalization to one standard space (MNI152NLin2009cAsym) was performed through nonlinear registration with antsRegistration (ANTs 2.3.3), using brain-extracted versions of both T1w reference and the T1w template. The following template was selected for spatial normalization: ICBM 152 Nonlinear Asymmetrical template version 2009c (Fonov et al., 2009, RRID:SCR_008796; TemplateFlow ID: MNI152NLin2009cAsym).

##### Functional Data Preprocessing

For each of the 3 BOLD blocks per participant, the following preprocessing was performed. First, a reference volume and its skull-stripped version were generated by aligning and averaging 1 single-band references (SBRefs). Head-motion parameters with respect to the BOLD reference (transformation matrices, and six corresponding rotation and translation parameters) are estimated before any spatiotemporal filtering using mcflirt (FSL 6.0.5.1:57b01774, Jenkinson et al., 2002). BOLD blocks were slice-time corrected to 0.696s (0.5 of slice acquisition range 0s-1.39s) using 3dTshift from AFNI (Cox & Hyde, 1997, RRID:SCR_005927). The BOLD time-series (including slice-timing correction when applied) were resampled onto their original, native space by applying the transforms to correct for head-motion. These resampled BOLD time-series will be referred to as preprocessed BOLD in original space, or just preprocessed BOLD. The BOLD reference was then co-registered to the T1w reference using bbregister (FreeSurfer) which implements boundary-based registration (Greve & Fischl, 2009). Co-registration was configured with six degrees of freedom. First, a reference volume and its skull-stripped version were generated using a custom methodology of fMRIPrep. Several confounding time-series were calculated based on the preprocessed BOLD: framewise displacement (FD), DVARS and three region-wise global signals. FD was computed using two formulations following Power (absolute sum of relative motions, Power et al., 2014) and Jenkinson (relative root mean square displacement between affines, Jenkinson et al., 2002). FD and DVARS are calculated for each functional block, both using their implementations in Nipype (following the definitions by Power et al., 2014). The three global signals are extracted within the CSF, the WM, and the whole-brain masks. Additionally, a set of physiological regressors were extracted to allow for component-based noise correction (CompCor, Behzadi et al., 2007). Principal components are estimated after high-pass filtering the preprocessed BOLD time-series (using a discrete cosine filter with 128s cut-off) for the two CompCor variants: temporal (tCompCor) and anatomical (aCompCor). tCompCor components are then calculated from the top 2% variable voxels within the brain mask. For aCompCor, three probabilistic masks (CSF, WM and combined CSF+WM) are generated in anatomical space. The implementation differs from that of Behzadi et al. in that instead of eroding the masks by 2 pixels on BOLD space, the aCompCor masks are subtracted a mask of pixels that likely contain a volume fraction of GM. This mask is obtained by dilating a GM mask extracted from the FreeSurfer’s aseg segmentation, and it ensures components are not extracted from voxels containing a minimal fraction of GM. Finally, these masks are resampled into BOLD space and binarized by thresholding at 0.99 (as in the original implementation). Components are also calculated separately within the WM and CSF masks. For each CompCor decomposition, the k components with the largest singular values are retained, such that the retained components’ time series are sufficient to explain 50 percent of variance across the nuisance mask (CSF, WM, combined, or temporal). The remaining components are dropped from consideration. The head-motion estimates calculated in the correction step were also placed within the corresponding confounds file. The confound time series derived from head motion estimates and global signals were expanded with the inclusion of temporal derivatives and quadratic terms for each (Satterthwaite et al., 2013). Frames that exceeded a threshold of 0.5 mm FD or 1.5 standardised DVARS were annotated as motion outliers. The BOLD time-series were resampled into standard space, generating a preprocessed BOLD block in MNI152NLin2009cAsym space. First, a reference volume and its skull-stripped version were generated using a custom methodology of fMRIPrep. All resamplings can be performed with a single interpolation step by composing all the pertinent transformations (i.e. head-motion transform matrices, susceptibility distortion correction when available, and co-registrations to anatomical and output spaces). Gridded (volumetric) resamplings were performed using antsApplyTransforms (ANTs), configured with Lanczos interpolation to minimize the smoothing effects of other kernels (Lanczos, 1964). Non-gridded (surface) resamplings were performed using mri_vol2surf (FreeSurfer).

### Behavioral Analysis

#### Preprocessing

As a sanity check, we confirmed that all participants scored above chance-level (25%) in the memory test after the last block of learning, and recalled more than half of the learned words in the posttest.

Participants’ written responses to the pseudo-word meaning in the posttest were coded as (1) matching the synonym, (2) meaningless, (3) creative, unexpected answers (e.g., when one consider a mismatch pseudo-word “human-kla”, the opposite of human, to be “animal”), and (4) unexpected but incorrect answers (e.g., when one confused the meaning of different affix forms, mistook “warm-ran” as “warm-kla’ and reported the meaning to be “cold”, the opposite of warm). We excluded the unexpected cases in the analysis, which concerns 4.8% of the trials. As a result of the paired presentations in the priming task (e.g., “white-kla” is always followed by “black”), when participants indicated the pseudo-word to be meaningful, they usually provided the target synonym to be the inferred meaning (Kendall’s τ = .98, p < .001). Therefore, we used the second measure (i.e., the percentage of inferred meaning to match the synonym word) as an indicator for participants’ explicit inference, given that it provides more certainty than a binary choice. Based on the posttest, we excluded participants who failed to learn the structural affixation rules (N = 4, of which 2 were the same participants excluded due to MRI quality control). They were defined as those who consider more than half of the pseudo-words in the mismatch condition to be meaningful, or more than half of the pseudo-words in the congruent condition to be meaningless.

#### Statistical Analyses

We performed the statistical analyses of behavioral data using generalized linear mixed models with the *glmmTMB* package (Version 1.9.11, Brooks et al., 2017) in R (Version 4.1.0; R Core Team, 2017). For analysis of the probe trials and the posttest, we used experimental conditions (congruent vs. incongruent vs. mismatch) as the predictor. Participants and items were included as random effects, with conditions as random slope for participants. The significance of conditions was assessed using the Type II Wald Chi-square test. We used the *multcomp* package (Version 1.4.17, Hothorn et al., 2008) to perform multiple comparisons between the three experimental conditions.

### fMRI Analysis

fMRI data were analyzed using SPM12, the Matlab-based RSA toolbox (Nili et al., 2014, https://github.com/rsagroup/rsatoolbox_matlab) and custom scripts written in MATLAB R2022b (Mathworks Inc.; https://nl.mathworks.com/products/matlab.html).

#### Univariate Analysis

To identify the neural representations of the newly composed pseudo-word meanings, we exploited the phenomenon of fMRI adaptation (also termed as “repetition suppression”; Barron et al., 2016; Grill-spector et al., 2006). fMRI adaptation relies on the fact that neurons show suppression (or “adaptation”) in their response to repeated presentation of stimuli to which they are sensitive. This phenomenon is often used as a tool to infer representational overlap by assessing the relative suppression/adaptation between two different stimuli. To test the univariate fMRI adaptation effects, we implemented an event-related generalized linear model (GLM) in SPM 12. The GLM modeled both events of the prime and the target, and contained separate onset regressors for each of the four experimental conditions (i.e., congruent, incongruent, and 2 times counterbalanced mismatch conditions to ensure the same amount of trials per type of affixes). Furthermore, the GLM contained an onset regressor for the probe trials and a button press regressor as a regressor of no interest. All regressors were convolved with a canonical haemodynamic response function. Because of the sensitivity of the blood oxygen level-dependent signal to motion and physiological noise, we included in the GLM the framewise displacement, six rigid-body motion parameters (three translations and three rotation), six anatomical component-based noise correction components (aCompCorr) and all the cosine regressors estimated by fmriprep as confound regressors for denoising. Each block was modeled separately within the GLM.

We compared congruent versus incongruent prime-related activities to assess the computational processes as a function of sequential order rules, and congruent versus incongruent target-related activities for the compositional outcome. In addition, we had included in our preregistration a contrast between congruent/incongruent versus mismatch conditions for successful inference. However, given that our participants did not distinguish incongruent and mismatch conditions in their behavioral responses, it makes little sense to investigate this contrast. Instead, we compared congruent versus mismatch prime- and target-related activities as a control regardless of sequential order rules (Supplementary Material 3). The contrast images of all participants were then analyzed as a second-level random effects analysis. Our a priori hypothesis suggests the engagement of a medial prefrontal-hippocampal network for compositional generalization. To test this, we performed SVC using an anatomically defined ROI combining the hippocampal formation (incl. hippocampus, entorhinal cortex, subiculum), as well as a functionally defined medial prefrontal (mPFC) ROI. The hippocampal mask is defined using the Juelich histological atlas with a probabilistic threshold of 50%. The mPFC mask is defined based on previous study on conjunctive representation of building blocks in specific relational positions (t_29_ > 2.5; Figure 3B in Schwartenbeck et al., 2023). We considered our results significant if they survived family-wise error (FWE) correction at the cluster-level of P < 0.05 within these masks. To explore the engagement of key language areas, we performed additional SVC using two anatomically defined masks: the left inferior gyrus, IFG, and the left anterior temporal lobe (ATL). Both masks are defined using the Harvard-Oxford cortical structural atlas with a probabilistic threshold of 30% (Supplementary Material 5). Activations in other brain regions were only considered if they survived whole-brain cluster-level FWE correction at P<0.05. All statistical parametric maps visualized in the manuscript are thresholded at P<0.001 uncorrected and unmasked for illustration.

#### Multivariate Representational Similarity Analysis

To decode the representations of both the relational rules and the newly inferred word meanings at the time of prime, we adopted a multivariate RSA approach. In brief, RSA allows for the comparison of representational geometry in computational models and neural response patterns in the brain elicited by our stimuli (Kriegeskorte et al., 2008; Nili et al., 2014). By comparing the representational dissimilarity matrix predicted by the computational models of interest to those measured in the brain, RSA can therefore uncover representations of interest, such as abstract relational rules in the brain.

To capture the multivariate representation of word meaning and affixation rules, we ran two additional GLMs. The first GLM modeled events of the prime and contained separate onset regressors for each of the 30 compositional pseudo-words regardless of whether they were congruent or incongruent (e.g., a single onset regressor was used for both “white-kla” and “kla-white”). The second GLM again modeled the prime, but separately for the congruent and the incongruent conditions (e.g., one onset regressor was used for “white-kla” and another one for “kla-white”). Both GLMs contained one regressor for all the prime trials in the mismatch condition, one regressor for all the target words, one regressor for the probe trials, and one regressor for the button presses as regressors of no interest. All regressors were convolved with a canonical haemodynamic response function. Both GLMs included the same confound regressors as in the univariate analysis. Each block was modeled separately within the GLMs.

Subsequently, we computed the neural representational dissimilarity matrices (RDMs) for the 30 items at prime from the parameter estimate maps derived from the GLMs described above. Pairwise correlation distance (one minus Pearson correlation coefficient) was used as distance metric. RSA was performed whole-brain using a searchlight approach with a spherical searchlight (radius = 7 mm, approx. 180 voxels).

We constructed two model RDMs of interest to probe for the representation of compositional word meaning and the representation of abstract task rules. First, we constructed a “meaning” model using the embedding vectors for the 30 target words (e.g., “black” in “whitekla-black”) from a word embedding model (Mandera et al., 2017). Word embedding provides learned representations in a vector space for text where words that have similar meanings have similar representations. Mandera et al. (2017) evaluated several prediction-based models on their performance on a large behavioral dataset on semantic priming. Here we used the model with the best fit to the behavioral data, namely, a Continuous Bag of Words model (CBOW) trained on SONAR-500 text corpus (Oostdijk et al., 2013) and a corpus of movie subtitles. We calculated the pairwise Pearson correlation distance between the embeddings of all target words, and constructed a 30 *30 distance matrix for target meaning representations. In addition to the target meaning model, we also constructed a stem meaning model based on the stems of the 30 prime words (e.g., “white” in “whitekla-black”). Second, we constructed a “rule” model, for which we considered a 30*30 binary-coded distance matrix where the rules are either the same (e.g., “good-kla” and “short-kla”), or different (“good-kla” and “kla-dog”).

Within each searchlight sphere in each participant, we compared the model RDMs and the neural RDMs using Kendall’s rank correlation. Both the searchlight of the neural RDMs and the comparison with the model RDMs were performed using the Matlab-based RSA toolbox (Nili et al., 2014). The resulting correlation coefficients were then submitted to a one-sample t-test using SPM12 (i.e., contrasting the obtained correlation against zero). Effects were tested for statistical significance using cluster-inference with a cluster-defining threshold of p < 0.001 and whole-brain cluster-level FWE correction at p < 0.05.

We additionally explored two ROI-based RSA analyses, using the same hippocampal mask and the left IFG mask used in the univariate analysis. The same procedure was performed for each structural ROI instead of each searchlight sphere. First-level coefficients were then used in the group-level one-sample one-side t-test. Since the meaning model and the rule model are to some degree correlated (Kendall’s τ = .22, p < .001), we additionally used partial correlation for calculating the participant-level correlation between the model RDMs and the neural RDMs, in order to evaluate their unique contribution to the neural representations. To estimate the non-noise variance in the neural data, and thereby upper bounds of any model in explaining the neural data, we computed the noise ceiling employ a leave-one-participant-out approach: For each participant we computed the correlation of the neural RDM with the average neural RDM of all other participants; we then took the average correlation as the estimate of the noise ceiling.

As a sanity check of the RSA procedure, we additionally computed two visual model RDMs for target word representations. The two RDMs both reflect the visual similarity of the target words participants see on the screen. They were computed as (1) the Levenshtein distance calculated using the “stringdist” library (van der Loo, 2014) in R; and (2) the pixel-wise Euclidean distance between individual words presented on the screen. Expectedly, these two RDMs were highly correlated (Kendall’s τ = 0.57, p < .001).

## Acknowledgements

This research was supported by the Language in Interaction consortium (Gravitation Grant 024.001.006 funded by the Dutch Research Council). We would like to thank Anna Aumeistere, Anna Petukhova and Diede Booltink for their assistance in data collection.

## Data and Code Availability

All the research data (e.g., data files, analysis scripts) associated with the current paper are shared in the Donders repository (https://data.donders.ru.nl/).

## Supplementary Materials

### Supplementary Material 1: Full behavioral results

#### Learning

During a pre-scanning training phase, 30 healthy participants were exposed to pairs of compositional pseudo-words along with their meanings (Figure 1A, see Supplementary Material 2 for a full set of stimuli). Each of these compositional pseudo-words comprises a known stem (e.g., “good” in “good-kla”) and an unknown affix (e.g., “kla”). We manipulated the mapping of the meaning to the affix based on their sequential position: e.g., “-kla” as a suffix means “the opposite”, whereas “kla-” as a prefix means “young version”. These position-dependent affixation rules allowed participants to compose unique meanings based on different sequential combinations of the affixes with the stems. Crucially, while the participants could infer the affixation rules from the exemplars, these rules were never made explicit to them. After four blocks of repetition, all participants were able to recall the meanings of the pseudo-words, evidenced by ceiling level performance on a subsequent memory task (mean_accuracy_ = 98.2 %, SD = 3.1%, Figure 1B).

#### Testing (in the scanner)

To test participants’ knowledge of the abstract structure rules, we presented them with a new set of compositional pseudo-words they had never encountered before (e.g. “white-kla” and “kla-white”) while they were in the scanner, and asked them to imagine the meanings of the words. These novel words could be either congruent with the sequential order rule they had learned (e.g. for “white-kla”, “-kla” as suffix means “the opposite”, and the opposite of white is “black”), or incongruent (e.g. for “kla-white”, “kla-” as a prefix means “the young version of”, while it is much more difficult to infer the meaning of the young version of white). We employed an fMRI adaptation paradigm in which the pseudo-words (“primes”) were presented in pairs with their synonym (“targets”) matched to the congruent condition (e.g., “black” was presented after “white-kla” and also after “kla-white”, Figure 1C). After 10% of the targets, participants were presented a probe question about whether the meaning of the target word was the same as that of the preceding prime (i.e., the compositional pseudo-word). This manipulation allowed us to assess whether participants successfully inferred the meaning of the novel compositional words using the relational structure rules. Analysis of performance on these probe catch trials revealed significantly higher probability of meaning-match responses in congruent (mean = 90.7%, SD = 16.5%) than incongruent trials (mean = 23.8%, SD = 32.3%; β = 4.52, SE = 0.78, z = 5.80, p < .001; Figure 1D). This evidenced reliance on the relational structure rules for inference. As a control condition, we also included mismatched pseudo-words as primes, where the stems were combined with alternative affixes, so that the meaning of the compositional pseudo-words did not match the target synonyms regardless of the position of the affix (e.g., ran-white = the color of white ≠ black; white-ran = the extreme version of white ≠ black). Note that in most of the cases, these words in the mismatch condition were also meaningless. As expected, participants did not consider these pseudo-words to match the meaning of the synonym (mean = 6.1%, SD = 7.8%). The degree to which the participants considered the novel word to match the synonym or not in the mismatch condition differed significantly from the congruent condition (β = 5.78, SE = 0.60, z = 9.60, p < .001), but not from the incongruent condition (β = 1.25, SE = 0.55, z = 2.27, p = .058; main effect between all three conditions: Χ^2^(2) = 93.51, p < .001).

#### Posttest

During the posttest, we asked participants explicitly to indicate whether they considered the novel pseudo-words that they had seen during the preceding MRI scan to be meaningful or not (Figure 1E). The pattern of the posttest results validated the probe results from the scanning session: The probability of “yes” responses to this question was significantly higher for the congruent pseudo-words than for the incongruent/mismatch pseudo-words (Figure 1F; mean_cong_ = 90.5%, SD_cong_ = 8.3%; mean_incong_ = 13.4%, SD_incong_ = 28.1%; mean_mismatch_ = 1.3%, SD_mismatch_ = 2.8%; Χ^2^(2) = 181.13, p < .001; congruent vs incongruent: β = 6.65, SE = 1.18, z = 5.64, p < .001; congruent vs. mismatch: β = 7.46, SE = 0.56, z = 13.43, p < .001; incongruent vs mismatch: β = 0.81, SE = 1.11, z = 0.73, p = .735).

### Supplementary Material 2: Experimental Stimuli

#### Supplementary Material 2A: Stimuli for Learning

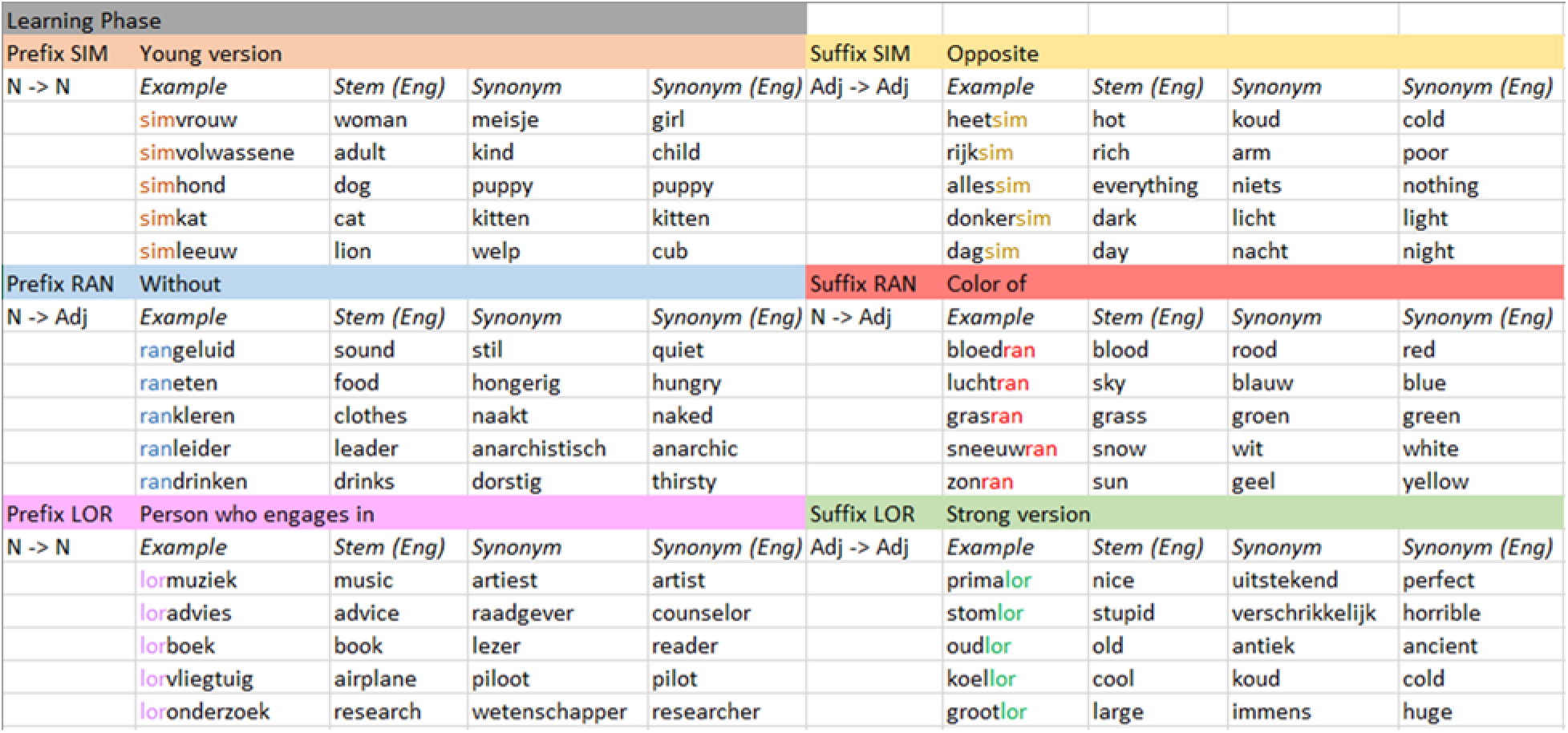

#### Supplementary Material 2B: Stimuli for Testing

| Testing Phase |  |  |  |  |  |  |  |  |  |
| --- | --- | --- | --- | --- | --- | --- | --- | --- | --- |
| Prefix SIM | Young version |  |  |  | Suffix SIM | Opposite |  |  |  |
| N -> N | Example | Stem (Eng) | Synonym | Synonym (Eng) | Adj -> Adj | Example | Stem (Eng) | Synonym | Synonym (Eng) |
|  | simpaard | horse | veulen | foal |  | kortsim | short | lang | long |
|  | simkip | chicken | kuiken | chick |  | jongsim | young | oud | old |
|  | simkoe | cow | kalf | calf |  | linkssim | left | rechts | right |
|  | simschaap | sheep | lam | lamb |  | volsim | full | leeg | empty |
|  | simmens | human | baby | baby |  | hoogsim | high | laag | low |
| Prefix RAN | Without |  |  |  | Suffix RAN | Color of |  |  |  |
| N -> Adj | Example | Stem (Eng) | Synonym | Synonym (Eng) | N -> Adj | Example | Stem (Eng) | Synonym | Synonym (Eng) |
|  | ranhaar | hair | kaal | bald |  | banaanran | banana | geel | yellow |
|  | ranzicht | sight | blind | blind |  | pupilran | pupil | zwart | black |
|  | ranreligie | religion | atheïstisch | atheistic |  | olifantran | elephant | grijs | grey |
|  | rankosten | expenses | gratis | free |  | aubergineran | eggplant | paars | purple |
|  | ranenergie | energy | moe | tired |  | mandarijnran | tangerine | oranje | orange |
| Prefix LOR | Person who engages in |  |  |  | Suffix LOR | Strong version |  |  |  |
| N -> N | Example | Stem (Eng) | Synonym | Synonym (Eng) | Adj -> Adj | Example | Stem (Eng) | Synonym | Synonym (Eng) |
|  | lorkeuken | kitchen | kok | chef |  | goed<sup>lor</sup> | good | geweldig | amazing |
|  | lorlied | song | zanger | singer |  | soms<sup>lor</sup> | sometimes | zelden | rarely |
|  | lortoneel | theatre | acteur | actor |  | warm<sup>lor</sup> | warm | heet | hot |
|  | lortrein | train | conducteur | train driver |  | klein<sup>lor</sup> | small | miniem | tiny |
|  | lorles | lesson | leerling | student |  | lastig<sup>lor</sup> | difficult | moeilijk | difficult |

Note: These six affix meanings were assigned to three affix forms (as prefix and suffix, respectively), counterbalanced across participants in six lists. In this way, we ensured that the priming effects were not confounded by visual input.

| pre | post | pre | post | pre | post |
| --- | --- | --- | --- | --- | --- |

| Young<br>version | opposite | without | color of | person<br>engaged | stronger<br>version |  |
| --- | --- | --- | --- | --- | --- | --- |
| SIM | SIM | RAN | RAN | LOR | LOR | list1 |
| SIM | SIM | LOR | LOR | RAN | RAN | list2 |
| RAN | RAN | SIM | SIM | LOR | LOR | list3 |
| RAN | RAN | LOR | LOR | SIM | SIM | list4 |
| LOR | LOR | SIM | SIM | RAN | RAN | list5 |
| LOR | LOR | RAN | RAN | SIM | SIM | list6 |

### Supplementary Material 3: Additional Univariate Analyses

#### Supplementary Material 3A: Full statistical outcome from the two main fMRI analyses

**Table S3-1.**
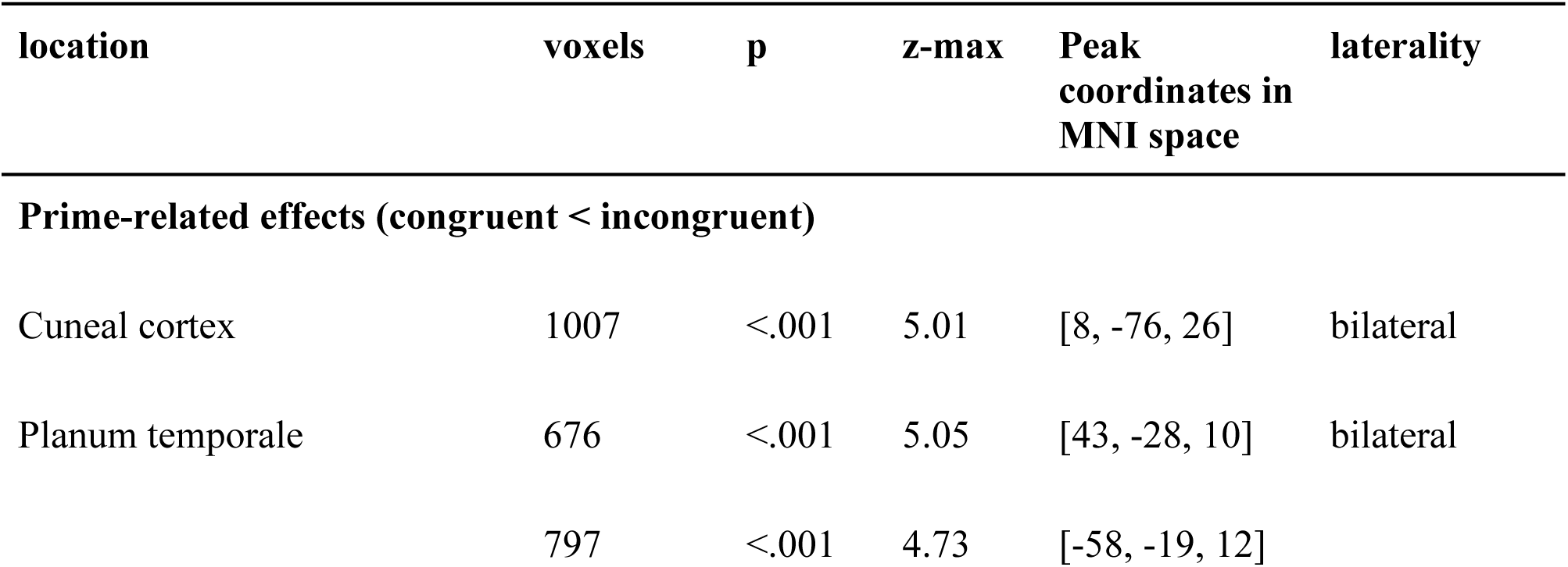
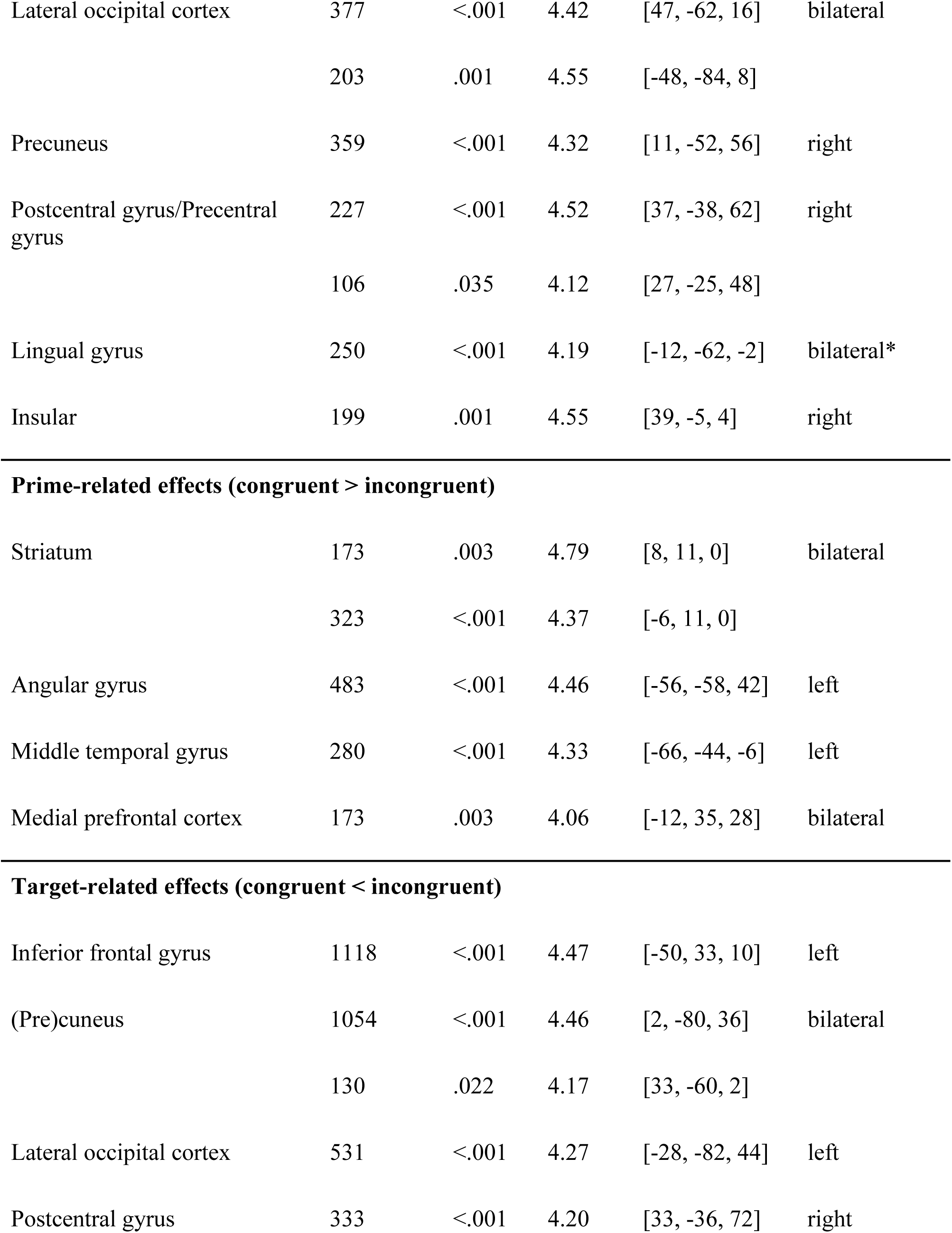
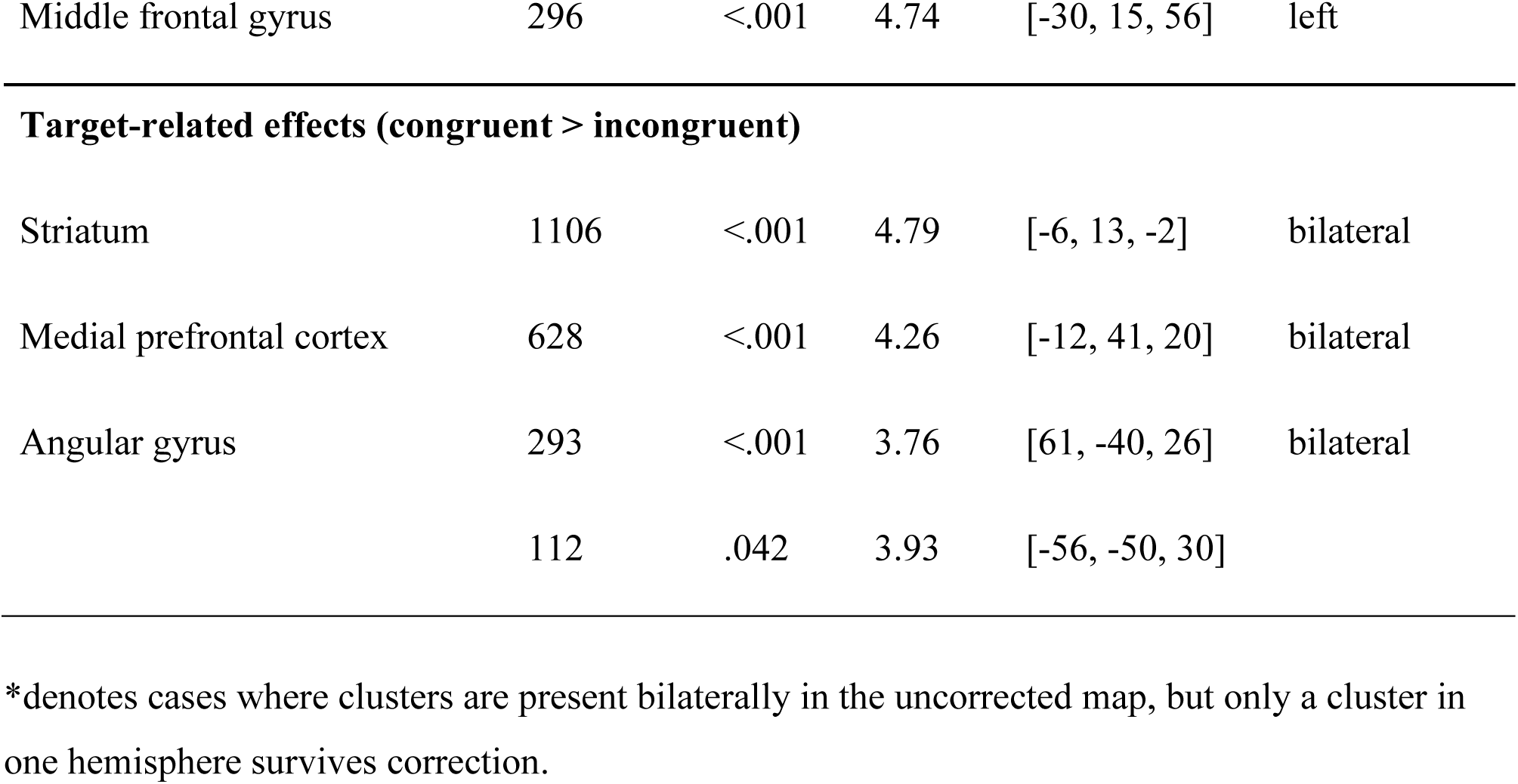
Univariate fMRI analysis of prime-related and target-related BOLD effects, as a function of congruent versus incongruent condition. Only significant clusters are included (p < .05, family-wise error corrected at the whole brain level). An uncorrected threshold of p < .001 is used for cluster forming. Results from the ROIs are reported in the main text.

| location | voxels | p | z-max | Peak<br>coordinates in<br>MNI space | laterality |
| --- | --- | --- | --- | --- | --- |
| **Prime-related effects (congruent < incongruent)** |  |  |  |  |  |
| Cuneal cortex | 1007 | <.001 | 5.01 | [8, -76, 26] | bilateral |
| Planum temporale | 676 | <.001 | 5.05 | [43, -28, 10] | bilateral |
|  | 797 | <.001 | 4.73 | [-58, -19, 12] |  |
| Lateral occipital cortex | 377 | <.001 | 4.42 | [47, -62, 16] | bilateral |
|  | 203 | .001 | 4.55 | [-48, -84, 8] |  |
| Precuneus | 359 | <.001 | 4.32 | [11, -52, 56] | right |
| Postcentral gyrus/Precentral gyrus | 227 | <.001 | 4.52 | [37, -38, 62] | right |
|  | 106 | .035 | 4.12 | [27, -25, 48] |  |
| Lingual gyrus | 250 | <.001 | 4.19 | [-12, -62, -2] | bilateral* |
| Insular | 199 | .001 | 4.55 | [39, -5, 4] | right |
| **Prime-related effects (congruent > incongruent)** |  |  |  |  |  |
| Striatum | 173 | .003 | 4.79 | [8, 11, 0] | bilateral |
|  | 323 | <.001 | 4.37 | [-6, 11, 0] |  |
| Angular gyrus | 483 | <.001 | 4.46 | [-56, -58, 42] | left |
| Middle temporal gyrus | 280 | <.001 | 4.33 | [-66, -44, -6] | left |
| Medial prefrontal cortex | 173 | .003 | 4.06 | [-12, 35, 28] | bilateral |
| **Target-related effects (congruent < incongruent)** |  |  |  |  |  |
| Inferior frontal gyrus | 1118 | <.001 | 4.47 | [-50, 33, 10] | left |
| (Pre)cuneus | 1054 | <.001 | 4.46 | [2, -80, 36] | bilateral |
|  | 130 | .022 | 4.17 | [33, -60, 2] |  |
| Lateral occipital cortex | 531 | <.001 | 4.27 | [-28, -82, 44] | left |
| Postcentral gyrus | 333 | <.001 | 4.20 | [33, -36, 72] | right |
| Middle frontal gyrus | 296 | <.001 | 4.74 | [-30, 15, 56] | left |
| **Target-related effects (congruent > incongruent)** |  |  |  |  |  |
| Striatum | 1106 | <.001 | 4.79 | [-6, 13, -2] | bilateral |
| Medial prefrontal cortex | 628 | <.001 | 4.26 | [-12, 41, 20] | bilateral |
| Angular gyrus | 293 | <.001 | 3.76 | [61, -40, 26] | bilateral |
|  | 112 | .042 | 3.93 | [-56, -50, 30] |  |

\*denotes cases where clusters are present bilaterally in the uncorrected map, but only a cluster in one hemisphere survives correction.

**Table S2-2.**
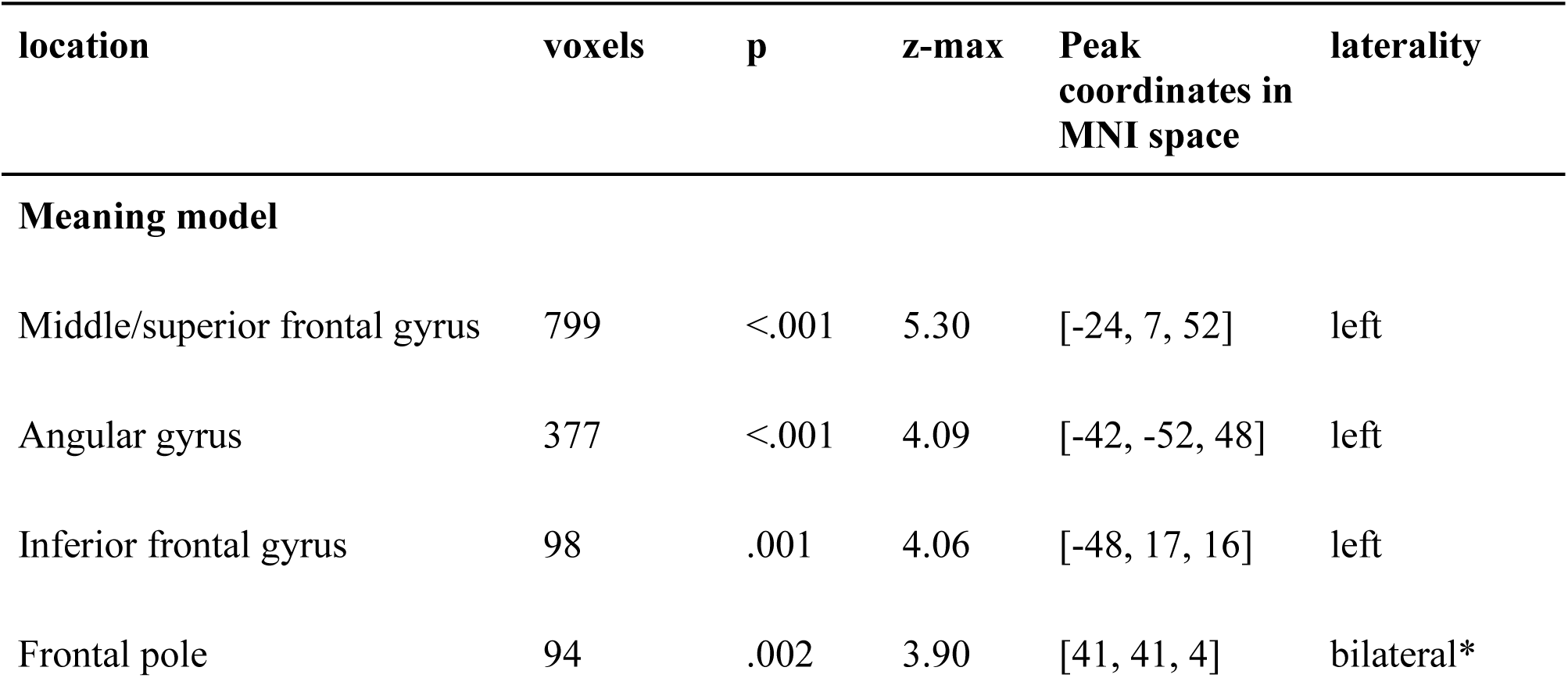
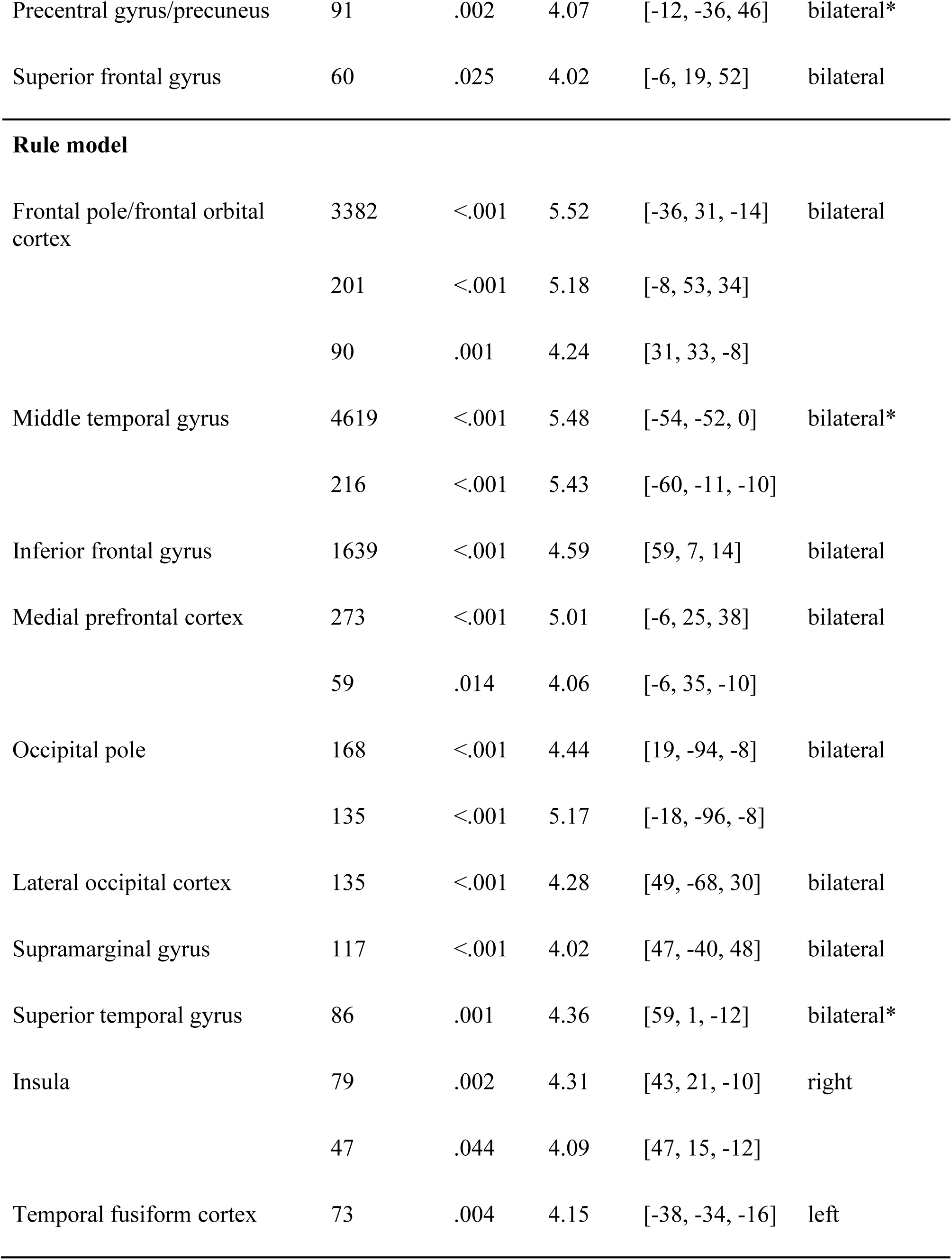
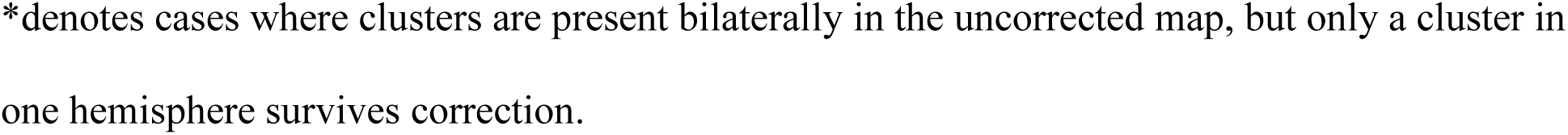
Multivariate fMRI representational similarity analysis of prime-related BOLD signals, compared with the meaning model and the rule model, respectively. Only significant clusters are included (p < .05, family-wise error corrected at the whole brain level). An uncorrected threshold of p < .001 is used for cluster forming.

| location | voxels | p | z-max | Peak coordinates in MNI space | laterality |
| --- | --- | --- | --- | --- | --- |
| **Meaning model** |  |  |  |  |  |
| Middle/superior frontal gyrus | 799 | <.001 | 5.30 | [-24, 7, 52] | left |
| Angular gyrus | 377 | <.001 | 4.09 | [-42, -52, 48] | left |
| Inferior frontal gyrus | 98 | .001 | 4.06 | [-48, 17, 16] | left |
| Frontal pole | 94 | .002 | 3.90 | [41, 41, 4] | bilateral* |
| Precentral gyrus/precuneus | 91 | .002 | 4.07 | [-12, -36, 46] | bilateral* |
| Superior frontal gyrus | 60 | .025 | 4.02 | [-6, 19, 52] | bilateral |
| **Rule model** |  |  |  |  |  |
| Frontal pole/frontal orbital cortex | 3382 | <.001 | 5.52 | [-36, 31, -14] | bilateral |
|  | 201 | <.001 | 5.18 | [-8, 53, 34] |  |
|  | 90 | .001 | 4.24 | [31, 33, -8] |  |
| Middle temporal gyrus | 4619 | <.001 | 5.48 | [-54, -52, 0] | bilateral* |
|  | 216 | <.001 | 5.43 | [-60, -11, -10] |  |
| Inferior frontal gyrus | 1639 | <.001 | 4.59 | [59, 7, 14] | bilateral |
| Medial prefrontal cortex | 273 | <.001 | 5.01 | [-6, 25, 38] | bilateral |
|  | 59 | .014 | 4.06 | [-6, 35, -10] |  |
| Occipital pole | 168 | <.001 | 4.44 | [19, -94, -8] | bilateral |
|  | 135 | <.001 | 5.17 | [-18, -96, -8] |  |
| Lateral occipital cortex | 135 | <.001 | 4.28 | [49, -68, 30] | bilateral |
| Supramarginal gyrus | 117 | <.001 | 4.02 | [47, -40, 48] | bilateral |
| Superior temporal gyrus | 86 | .001 | 4.36 | [59, 1, -12] | bilateral* |
| Insula | 79 | .002 | 4.31 | [43, 21, -10] | right |
|  | 47 | .044 | 4.09 | [47, 15, -12] |  |
| Temporal fusiform cortex | 73 | .004 | 4.15 | [-38, -34, -16] | left |

\*denotes cases where clusters are present bilaterally in the uncorrected map, but only a cluster in one hemisphere survives correction.

#### Supplementary Material 3B: Additional analysis using small volume correction

To probe the role of the language network in meaning inference, we additionally considered two anatomically defined masks, one for left anterior temporal lobe (left ATL) and one for left inferior frontal gyrus (left IFG; Supplementary Material 5). Apart from the above-mentioned fMRI adaptation effect in left IFG for incongruent versus congruent targets, there was no evidence for prime-related effect in the left IFG (no suprathreshold clusters found after SVC) nor target-related effect at left ATL (p_FWE_ = .082, K_E_ = 4, Z_max_ = 4.34, [−50, 1, −28], SVC). There was, however, greater activity in the left ATL during incongruent than congruent primes (p_FWE_ = .018, K_E_ = 22, Z_max_ = 3.71, [−62, −7, 2], SVC), which nevertheless merged with a bilateral temporal cluster and more likely reflected some general-purpose auditory processing (e.g., rehearsal of the composed meaning in order to make a comparison at target).

#### Supplementary Material 3C: Univariate Analysis of Congruent versus Mismatch Contrasts

##### Prime-related Activities

Comparison of mismatch versus congruent prime-related fMRI activity revealed greater activity in a multiple temporal and parietal areas, including the precuneus, the lateral occipital cortex, and the lingual gyrus (Figure S3A, Table S3). There was greater activity during mismatch than congruent primes in the hippocampal formation (p_FWE_ = .003, K_E_ = 84, Z_max_ = 4.19, [19, −9, −18], SVC), perhaps reflecting greater efforts to resolve the generalization-based composition challenge during mismatch than congruent primes. There was no evidence for any effect of prime type on activity in the mPFC or the left IFG (no suprathreshold clusters found after SVC). There was greater activity in the left ATL during mismatch than congruent primes (p_FWE_ = .034, K_E_ = 14, Z_max_ = 3.50, [−60, −9, 2], SVC), but merging with a bilateral temporal cluster. Interestingly, we also observe greater activation during congruent versus incongruent primes in the striatum (Figure S3C, Table S3).

##### Target-related Activities

Comparison of neural signals at the target word after a congruent versus mismatch compositional prime word revealed greater fMRI adaptation in a broad network of brain regions, including the (pre)cuneus, the postcentral gyrus, and the middle frontal gyrus (Figure S3; Table S3B). Critically, we also observed greater adaptation in the left inferior frontal cortex (Figure S3D). There was greater activity in the left ATL during mismatch than congruent target (p_FWE_ = .003, K_E_ = 63, Z_max_ = 4.54, [−62, −5, 2], SVC), but again merging with a bilateral temporal cluster.

There was no evidence for fMRI adaptation in the hippocampal formation during target words after congruent versus mismatch primes (no suprathreshold clusters found after SVC). Moreover, also in contrast to our hypothesis, activity in mPFC was actually greater at targets following congruent than mismatch primes. In addition, a similar pattern of greater activation during congruent versus mismatch targets was seen in the striatum.

**Figure S3.**
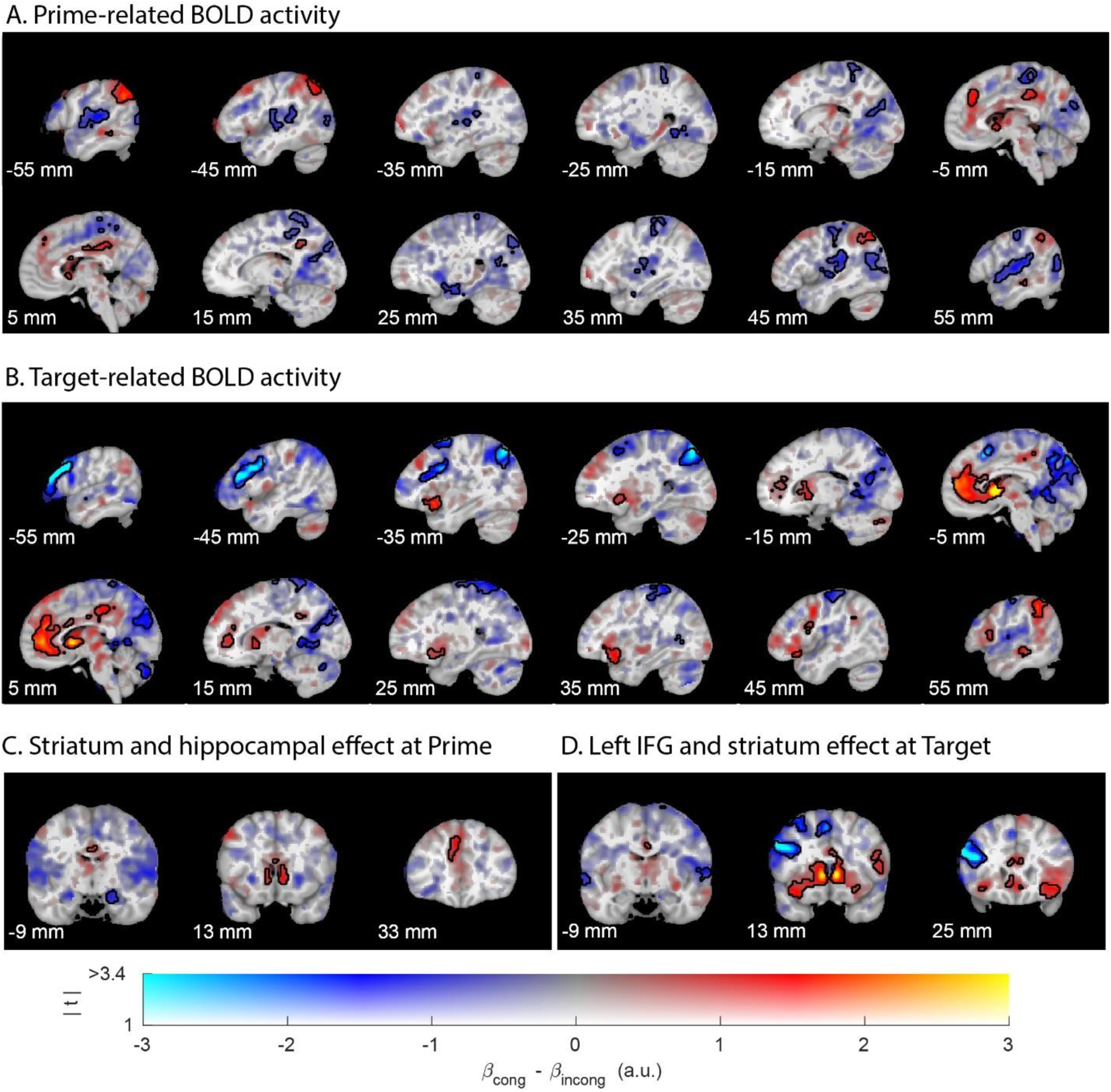
Univariate fMRI effects of novel compositional words meaning computation (prime-related activity) and representation (target-related activity). (A) fMRI effects of congruent versus mismatch prime-related BOLD activity. (B) fMRI effects of congruent versus mismatch target-related BOLD activity (in blue: fMRI adaptation). (C) Prime-related effects of interest. (D) Target-related effects of interest. The hue indexes the size of the parameter estimate, and the opacity indexes the unthresholded t values. Significant clusters (cluster-level corrected, FWE, p < .05) are encircled in solid contours. All coordinates are provided in the MNI space.

**Table S3.**
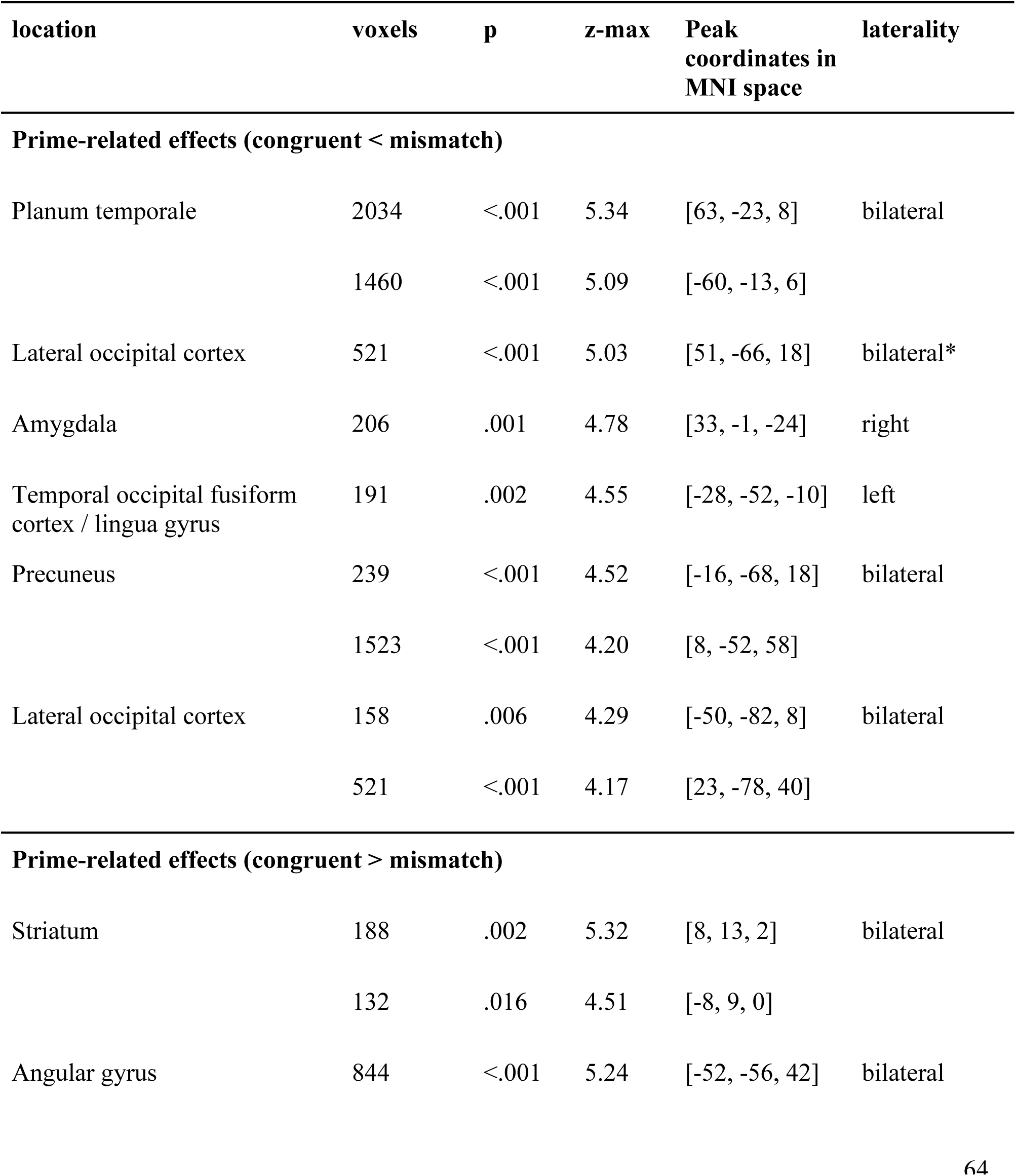
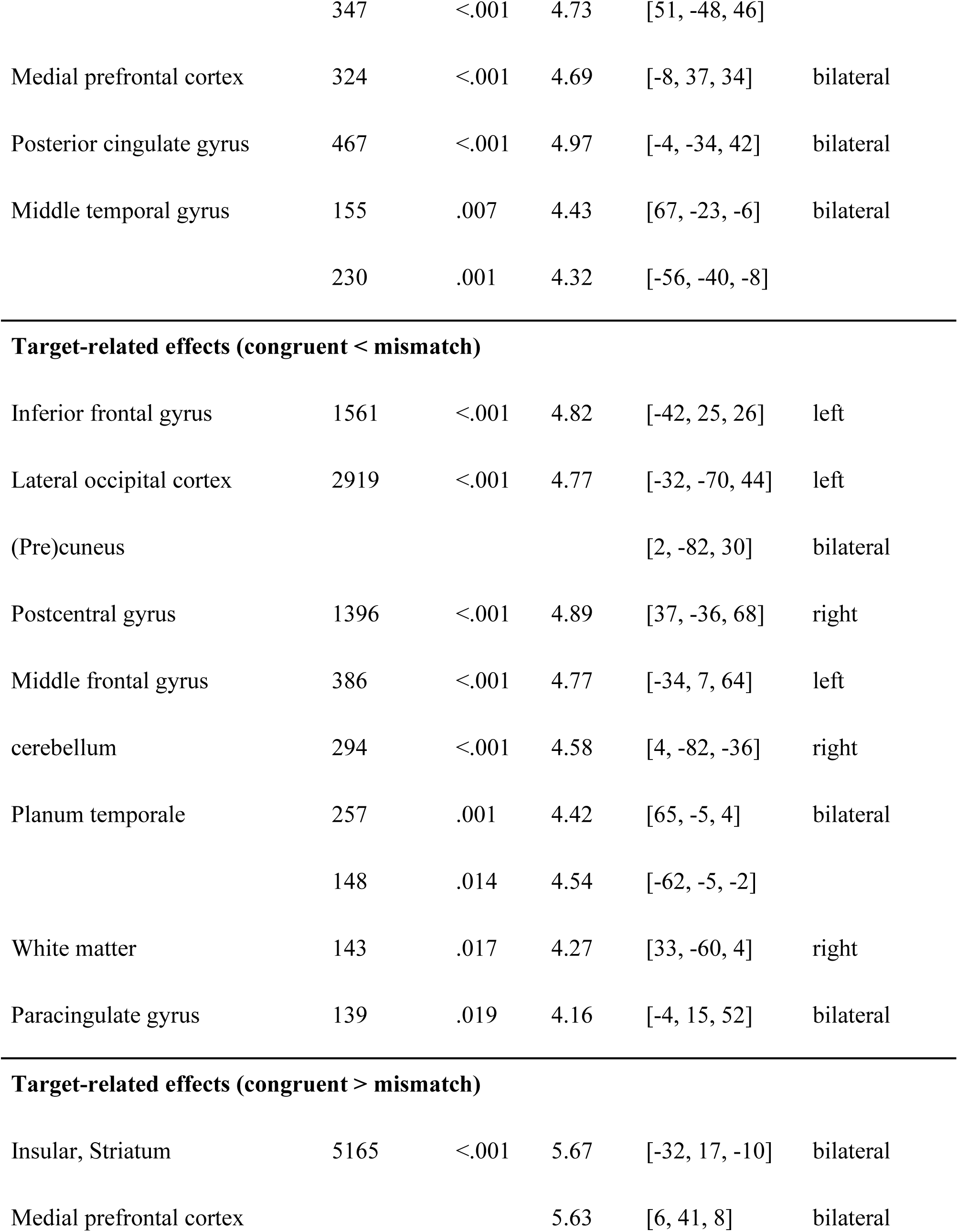
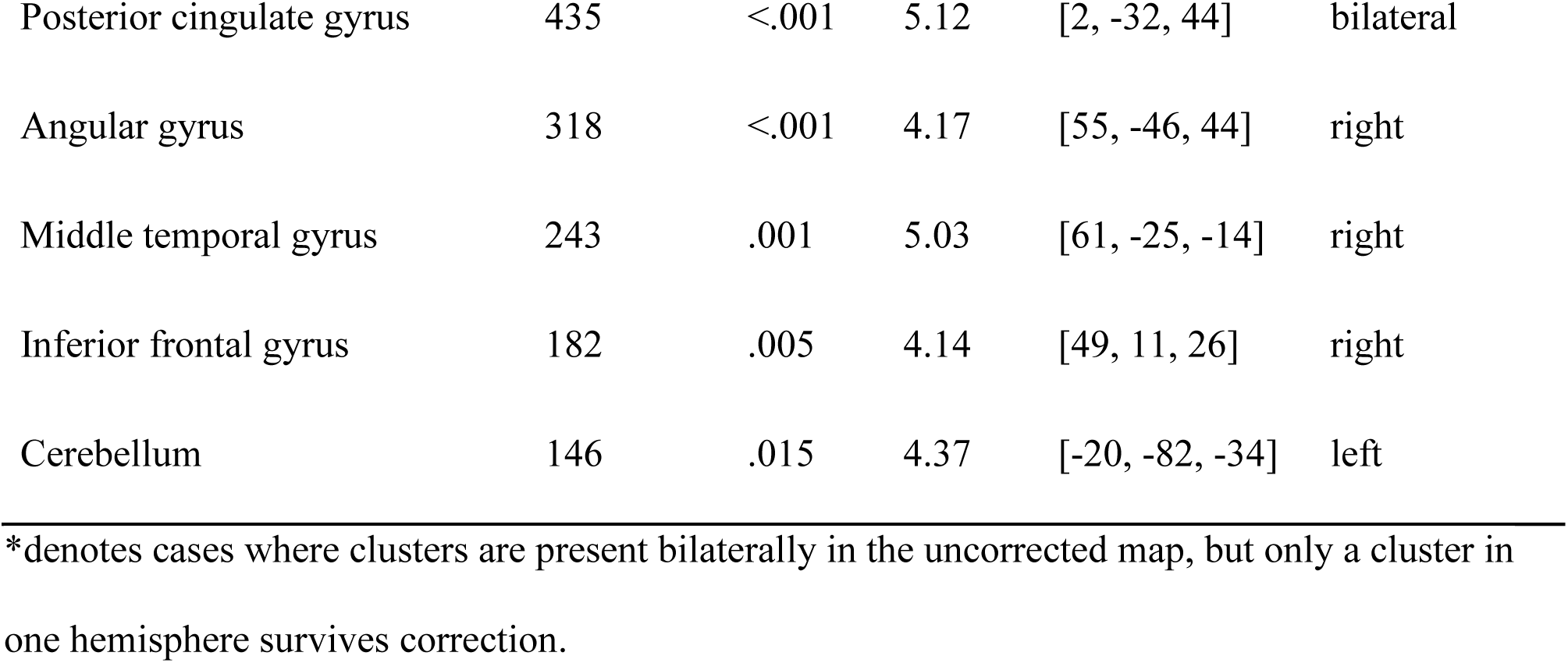
Univariate fMRI analysis of prime-related and target-related BOLD effects, as a function of congruent versus mismatch condition. Only significant clusters are included (p < .05, family-wise error corrected at the whole brain level). An uncorrected threshold of p < .001 is used for cluster forming. Results from the ROIs are reported below.

| location | voxels | p | z-max | Peak coordinates in MNI space | laterality |
| --- | --- | --- | --- | --- | --- |
| Prime-related effects (congruent < mismatch) |  |  |  |  |  |
| Planum temporale | 2034 | <.001 | 5.34 | [63, -23, 8] | bilateral |
|  | 1460 | <.001 | 5.09 | [-60, -13, 6] |  |
| Lateral occipital cortex | 521 | <.001 | 5.03 | [51, -66, 18] | bilateral* |
| Amygdala | 206 | .001 | 4.78 | [33, -1, -24] | right |
| Temporal occipital fusiform cortex / lingua gyrus | 191 | .002 | 4.55 | [-28, -52, -10] | left |
| Precuneus | 239 | <.001 | 4.52 | [-16, -68, 18] | bilateral |
|  | 1523 | <.001 | 4.20 | [8, -52, 58] |  |
| Lateral occipital cortex | 158 | .006 | 4.29 | [-50, -82, 8] | bilateral |
|  | 521 | <.001 | 4.17 | [23, -78, 40] |  |
| Prime-related effects (congruent > mismatch) |  |  |  |  |  |
| Striatum | 188 | .002 | 5.32 | [8, 13, 2] | bilateral |
|  | 132 | .016 | 4.51 | [-8, 9, 0] |  |
| Angular gyrus | 844 | <.001 | 5.24 | [-52, -56, 42] | bilateral |
|  | 347 | <.001 | 4.73 | [51, -48, 46] |  |
| Medial prefrontal cortex | 324 | <.001 | 4.69 | [-8, 37, 34] | bilateral |
| Posterior cingulate gyrus | 467 | <.001 | 4.97 | [-4, -34, 42] | bilateral |
| Middle temporal gyrus | 155 | .007 | 4.43 | [67, -23, -6] | bilateral |
|  | 230 | .001 | 4.32 | [-56, -40, -8] |  |
| **Target-related effects (congruent < mismatch)** |  |  |  |  |  |
| Inferior frontal gyrus | 1561 | <.001 | 4.82 | [-42, 25, 26] | left |
| Lateral occipital cortex | 2919 | <.001 | 4.77 | [-32, -70, 44] | left |
| (Pre)cuneus |  |  |  | [2, -82, 30] | bilateral |
| Postcentral gyrus | 1396 | <.001 | 4.89 | [37, -36, 68] | right |
| Middle frontal gyrus | 386 | <.001 | 4.77 | [-34, 7, 64] | left |
| cerebellum | 294 | <.001 | 4.58 | [4, -82, -36] | right |
| Planum temporale | 257 | .001 | 4.42 | [65, -5, 4] | bilateral |
|  | 148 | .014 | 4.54 | [-62, -5, -2] |  |
| White matter | 143 | .017 | 4.27 | [33, -60, 4] | right |
| Paracingulate gyrus | 139 | .019 | 4.16 | [-4, 15, 52] | bilateral |
| **Target-related effects (congruent > mismatch)** |  |  |  |  |  |
| Insular, Striatum | 5165 | <.001 | 5.67 | [-32, 17, -10] | bilateral |
| Medial prefrontal cortex |  |  | 5.63 | [6, 41, 8] | bilateral |
| Posterior cingulate gyrus | 435 | <.001 | 5.12 | [2, -32, 44] | bilateral |
| Angular gyrus | 318 | <.001 | 4.17 | [55, -46, 44] | right |
| Middle temporal gyrus | 243 | .001 | 5.03 | [61, -25, -14] | right |
| Inferior frontal gyrus | 182 | .005 | 4.14 | [49, 11, 26] | right |
| Cerebellum | 146 | .015 | 4.37 | [-20, -82, -34] | left |

\*denotes cases where clusters are present bilaterally in the uncorrected map, but only a cluster in one hemisphere survives correction.

### Supplementary Material 4: Additional RSA Analysis

#### Supplementary Material 4A: Visual RDM, Target-related activity

Before conducting the RSA of interest, we first validated our analysis approach by using two visual model RDMs to predict the target-related neural activity. These RDMs reflect the visual similarity of the target words participants saw on the screen. The RDMs were computed as (1) the Levenshtein distance calculated using the “stringdist” library (van der Loo, 2014) in R; and (2) the pixel-wise Euclidean distance between individual words presented on the screen. As expected, the two RDMs were highly correlated (Kendall’s τ = 0.57, p < .001). RSA results showed that the visual similarity between target words was reflected in visual cortical areas. Thus, this control analysis confirmed our analysis approach.

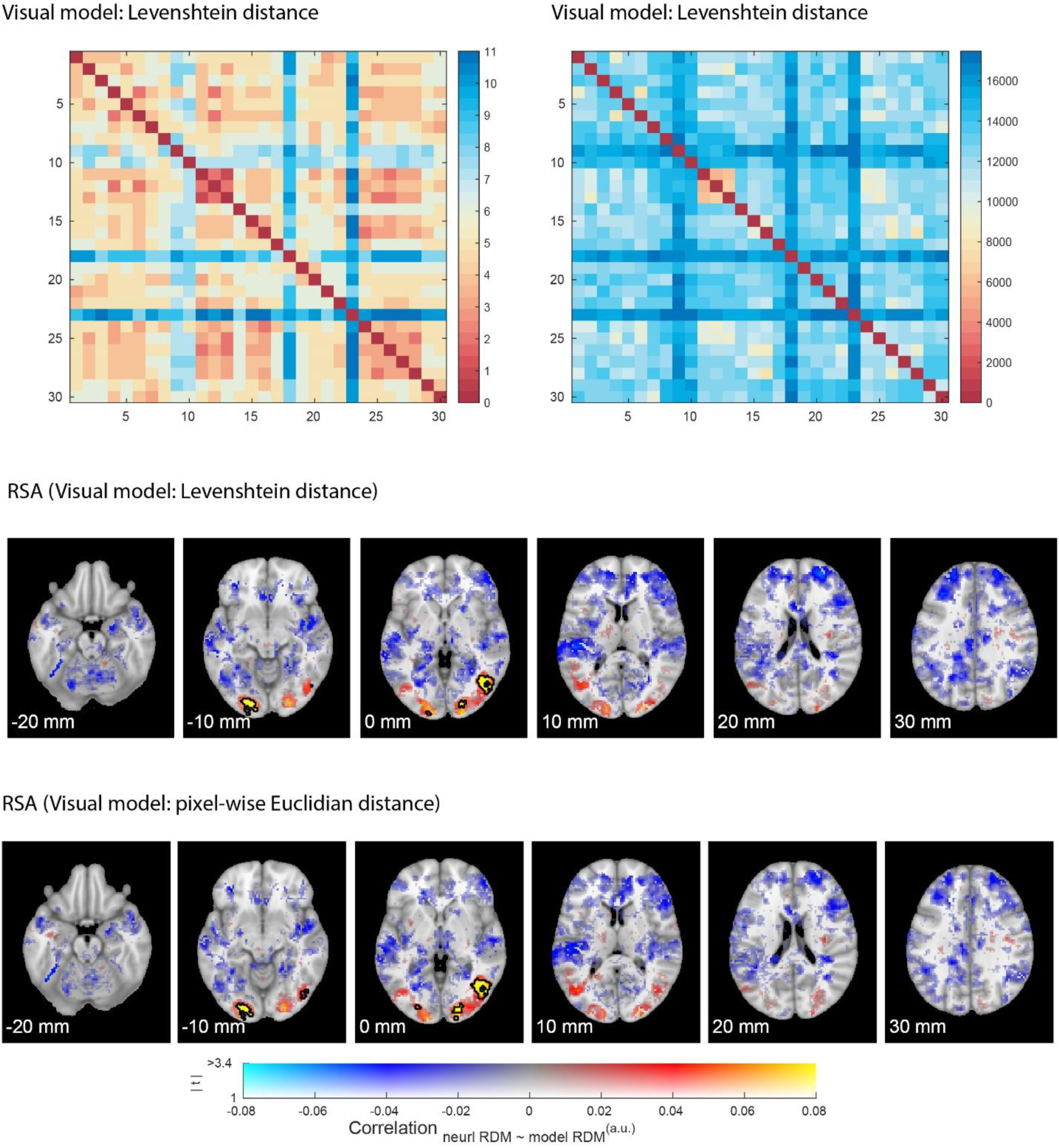

#### Supplementary Material 4B: Target meaning RDM, Target-related activity

Target-related neural representation was predicted by the same target-meaning model, reflected in a broad, left-lateralized network (incl. left IFC, left angular gyrus and left ATL).

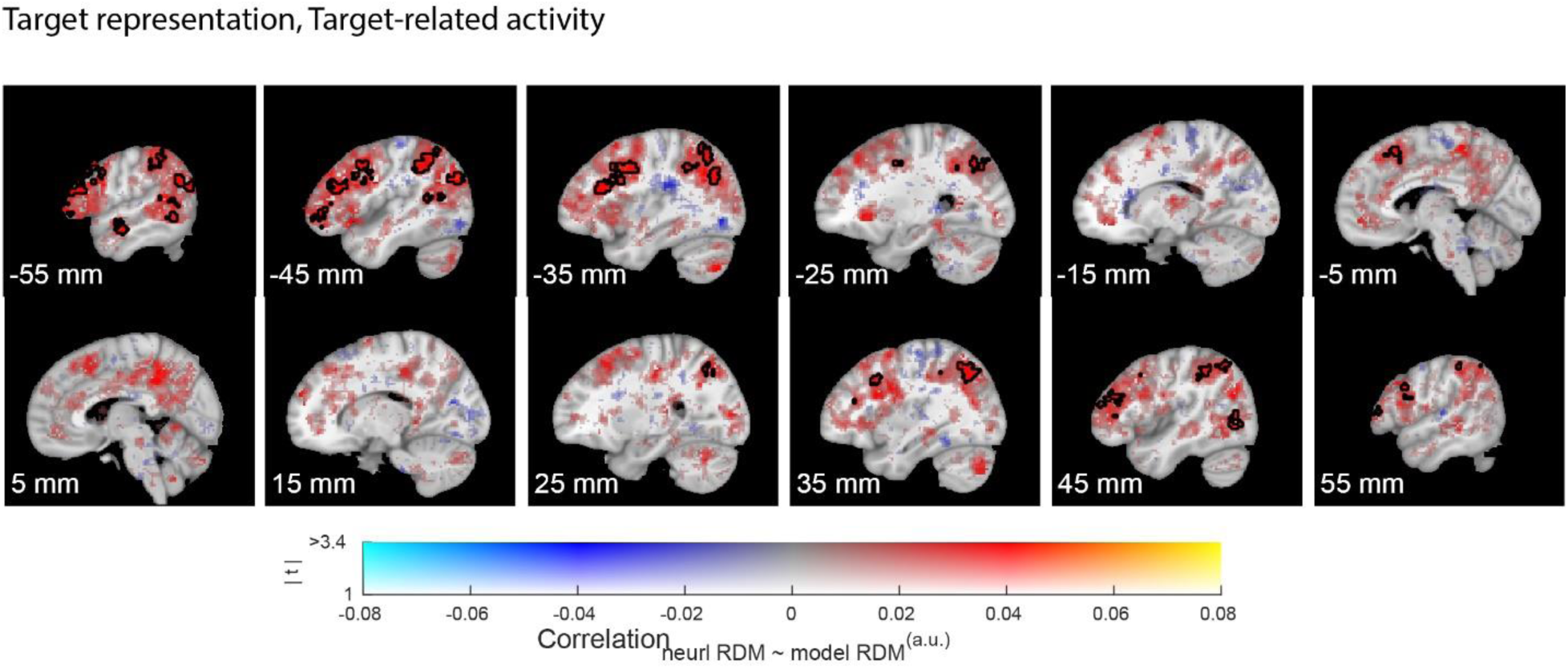

#### Supplementary Material 4C: Stem meaning RDM, Prime-related activity

Prime-related neural representation cannot be captured by an alternative meaning model which describes the similarities between stem meanings (e.g., “white” in “whitekla” is more similar to “light” in “lightkla”, compared to “sad” in “sadkla”; Kendall’s τ_stem-target_ = .13; all cluster-level ps > .841).

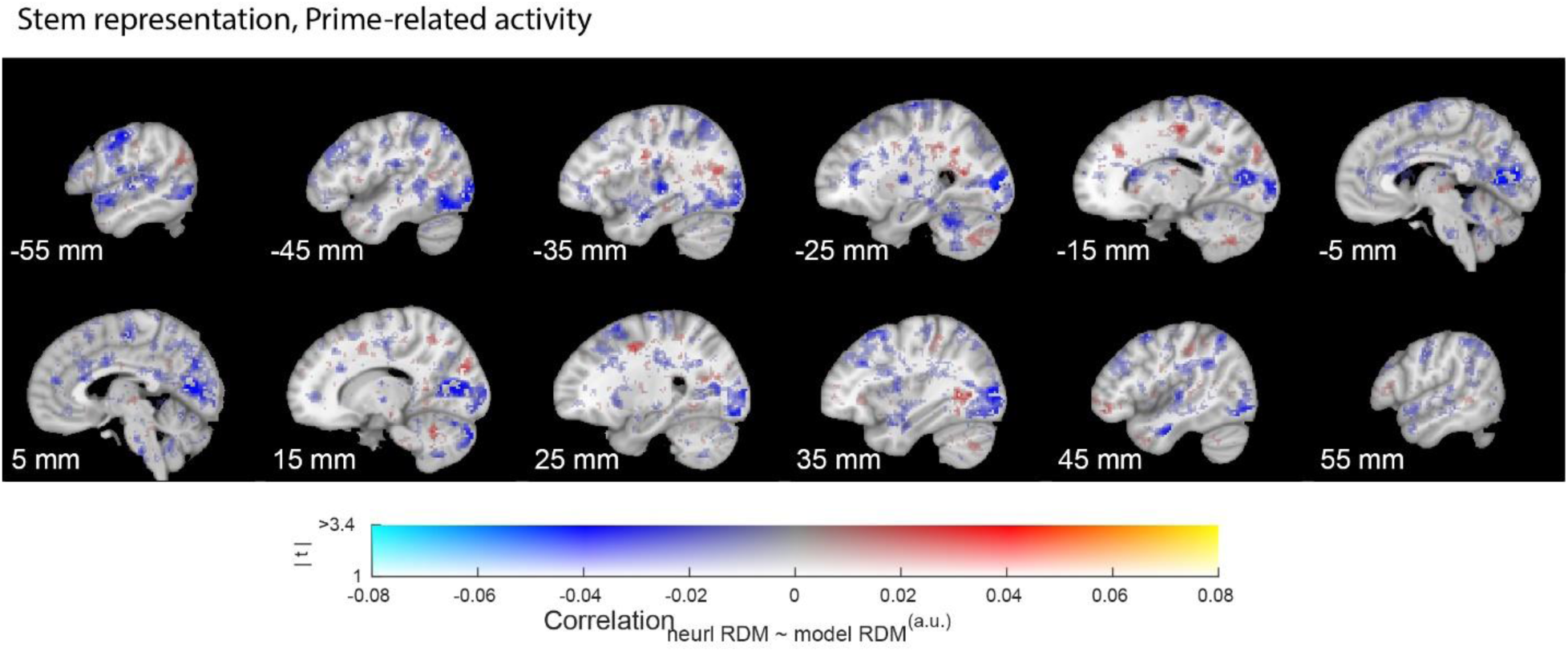

#### Supplementary Material 4D: Full ROI-based RSA

Based on our a priori hypothesis and the univariate analysis, we further explored two ROIs: the hippocampus and the left IFG, using the masks also used in the univariate analysis.

The ROI-based RSA (Figure 3E) confirmed the whole-brain results that left IFG represents both meanings (Mean_Kendall_τ_ = .04, SD = .06, t_29_ = 3.74, p < .001) and rules (Mean_Kendall_τ_ = .04, SD = .03; t_29_ = 7.70, p < .001). These results hold even after excluding the shared variance between the two models (meaning representation: Mean_Kendall_τ_partial_ = .04, SD = .07, t_29_ = 2.93, p = .003; rule representation: Mean_Kendall_τ_partial_ = .03, SD = .03, t_29_ = 5.24, p < .001; correlation between the two models: Kendall’s τ = .22, p < .001). In contrast, the hippocampal results were unconvincing considering the very low noise ceiling (meaning representation: Mean_Kendall_τ_ = .0003, SD = .04, t_29_ = 0.05, p = 0.482; rule representation: Mean_Kendall_τ_ = .01, SD = .03, t_29_ = 1.45, p = .079; noise ceiling = .004, calculated using a leave-one-out approach).

Furthermore, we explored the meaning representation and rule representation for congruent and incongruent words respectively (Figure 3E, right panel). We hypothesize that the meaning of a target word “black” is more likely to be inferred from a congruent prime “whitekla”, than from an incongruent prime “klawhite”. In contrast, the relational rules are likely recruited regardless of whether the compositional word is meaningful or not. This is, however, not confirmed by our ROI analysis of the left IFG: the target meaning representation can be decoded at the time of both congruent (Mean_Kendall_τ_partial_ = .02, SD = .05; t_29_ = 1.85, p = .037) and incongruent primes (Mean_Kendall_τ_partial_ = .04, SD = .05; t_29_ = 4.05, p < .001). While the effect was numerically larger for incongruent than congruent words, the difference was not statistically significant (t_29_ = −1.82, p = .080). On the contrary, rule representation is better captured in the congruent condition compared with the incongruent condition (congruent: Mean_Kendall_τ_partial_ = .03, SD = .03; t_29_ = 5.54, p < .001; incongruent: Mean_Kendall_τ_partial_ = .006, SD = .03; t_29_ = 1.47, p = .076; Paired t_29_ = 3.51, p = .001). The same pattern of differences between congruent and incongruent representations is confirmed in the whole-brain analysis (Supplementary Material 4E).

#### Supplementary Material 4E: Full brain RSA analysis of congruent and incongruent prime-related neural activities

Both the congruent and incongruent prime-related neural activities showed similar RSA outcome as the main analysis (i.e., when congruent and incongruent words were combined). There is no difference between the two conditions on the whole brain level for the meaning representation (Figure S4E-1, bottom panel, no suprathreshold clusters). In contrast, the congruent condition showed a stronger left frontal and temporal effect than the incongruent condition in rule representation (Figure S4E-2). These results confirmed the ROI-based RSA reported in the main text.

**Figure S4E-1.**
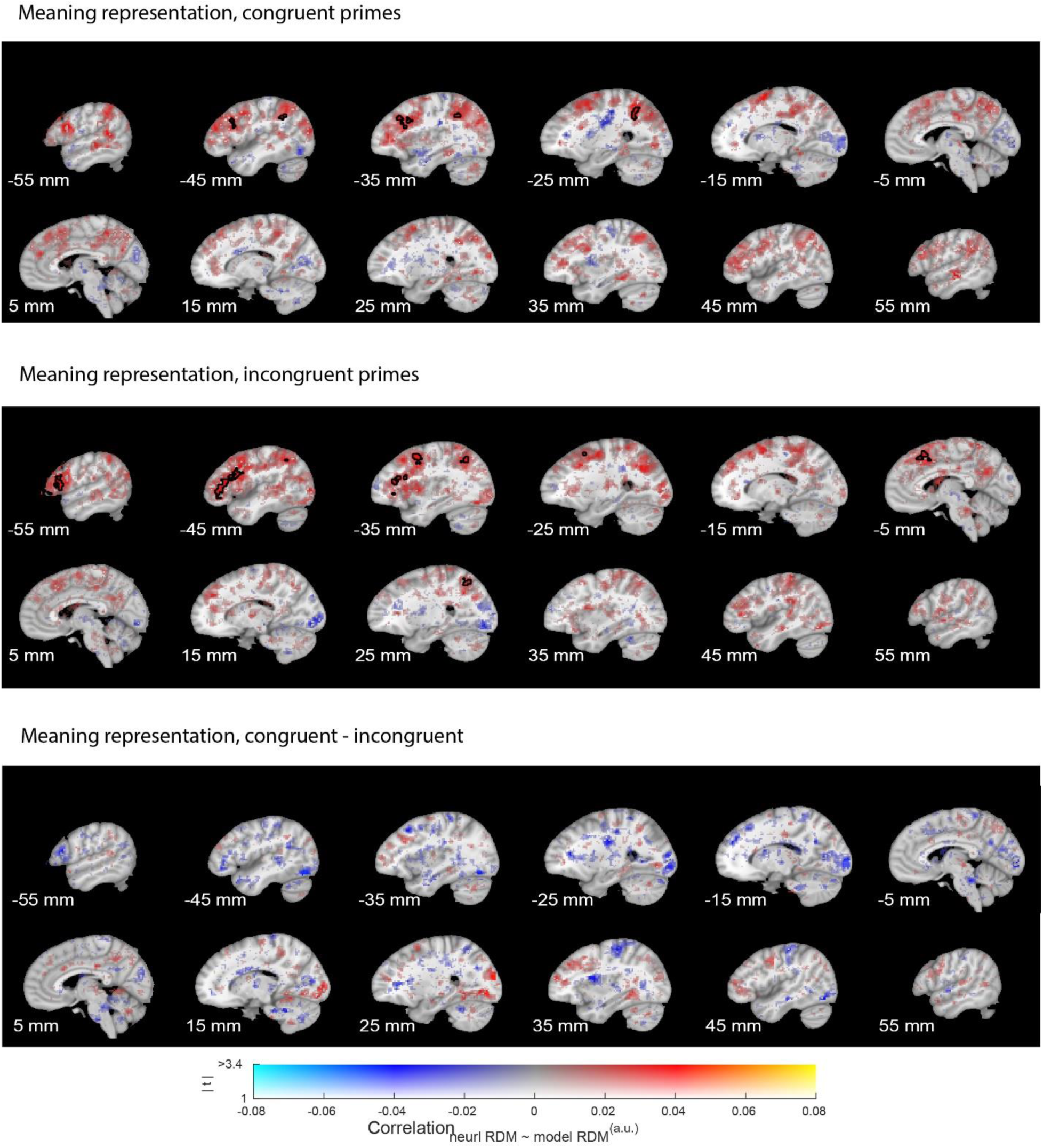
Whole-brain searchlight RSA outcome using the meaning model, performed for the congruent (top panel) and the incongruent (middle panel) conditions respectively, as well as their contrast (bottom panel).

**Figure S4E-2.**
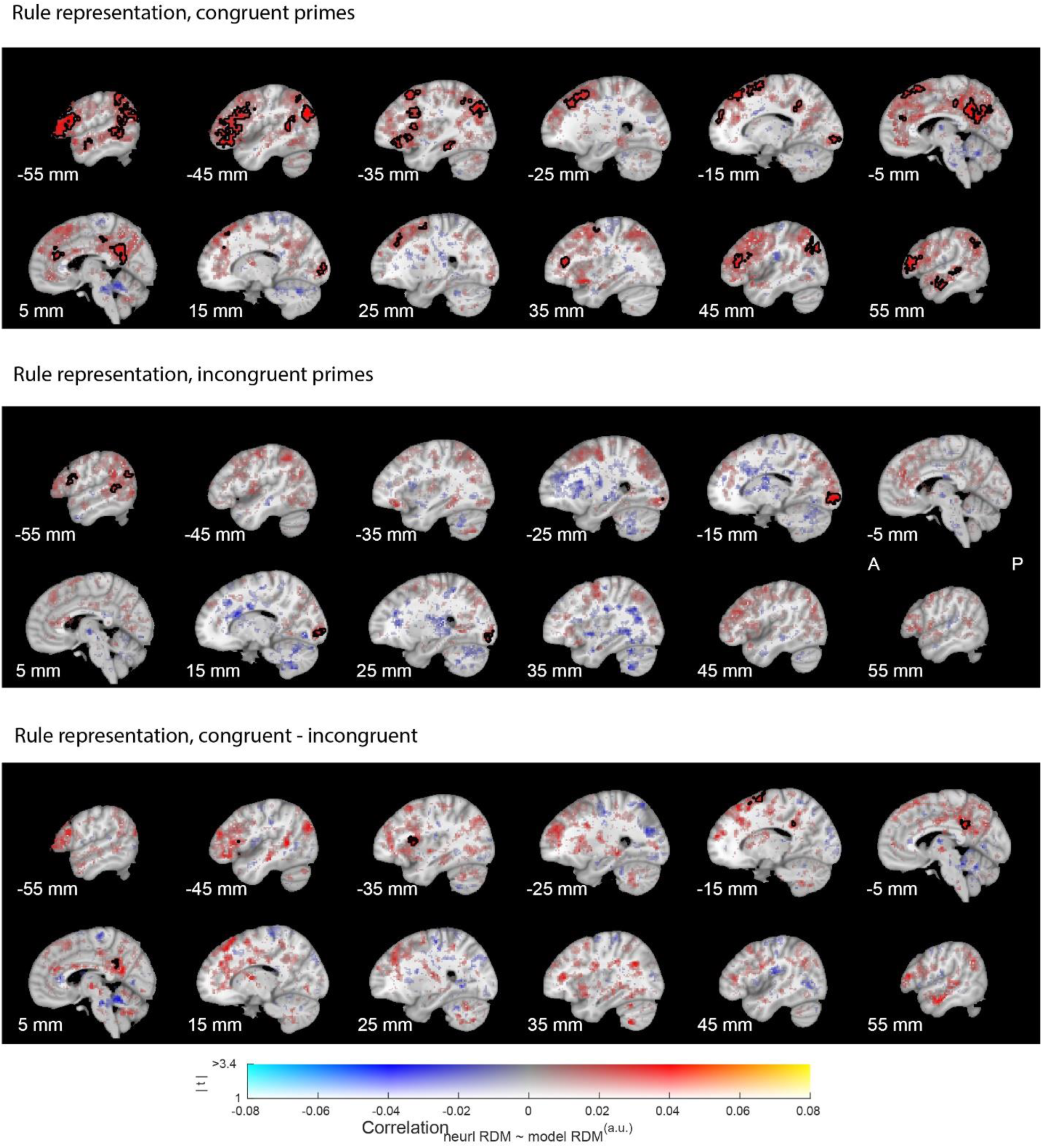
Whole-brain searchlight RSA outcome using the rule model, performed for the congruent (top panel) and the incongruent (middle panel) conditions respectively, as well as their contrast (bottom panel).

### Supplementary Material 5: Functional and Anatomical ROIs

**Figure S5.**
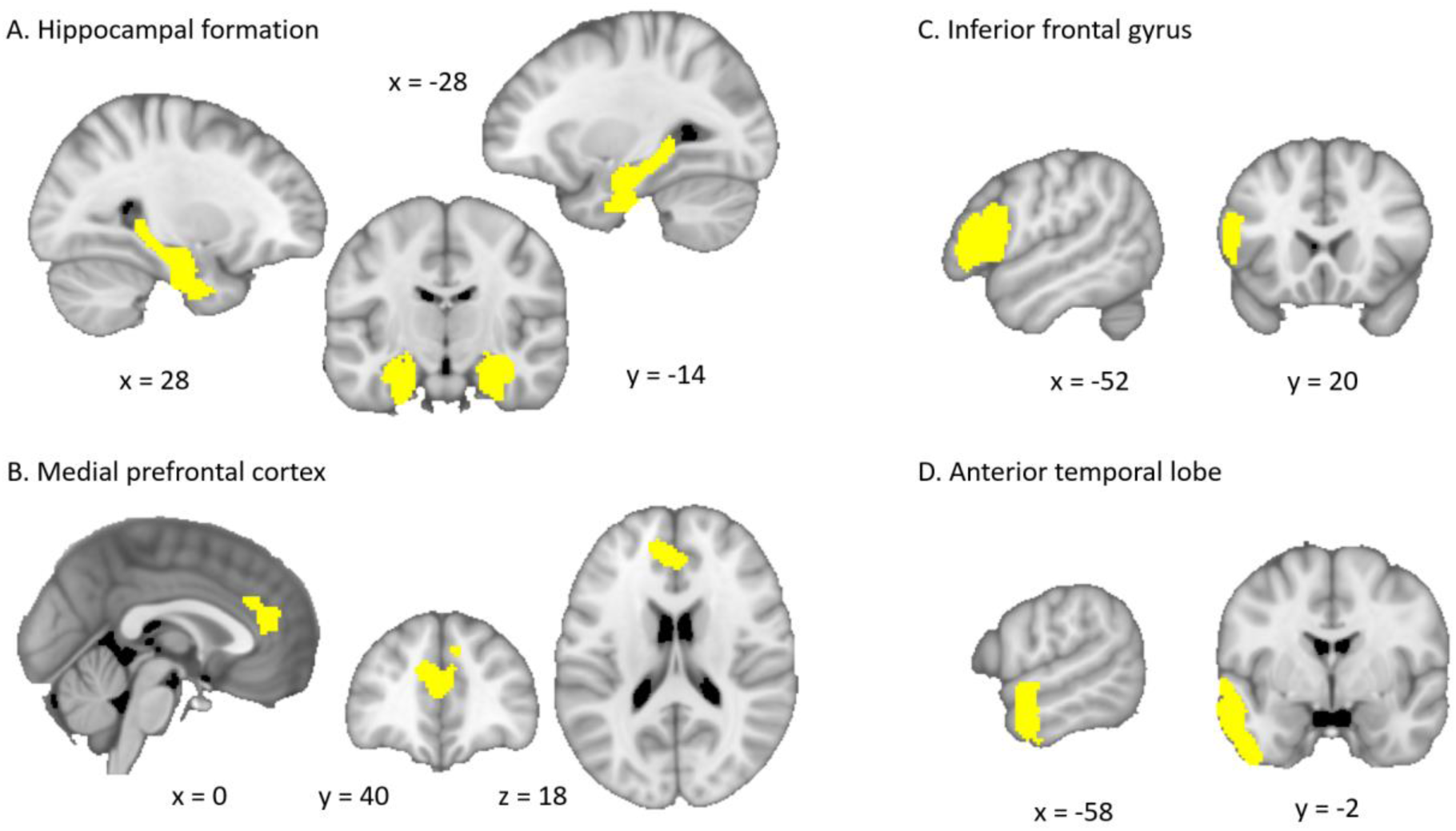
Functional and anatomical ROIs. (A) Anatomically defined ROI of the hippocampal formation, combining the hippocampal formation (incl. hippocampus, entorhinal cortex, subiculum). The hippocampal mask is defined using the Juelich histological atlas with a probabilistic threshold of 50%; (B) Functionally defined medial prefrontal (mPFC) ROI. The mPFC mask is defined based on previous study on conjunctive representation of building blocks in specific relational positions (t_29_ > 2.5; Figure 3B in Schwartenbeck et al., 2023). (C) Anatomically defined ROI consisting of the left inferior gyrus (IFG). (D) Anatomically defined ROI consisting of the left anterior temporal lobe (ATL). Both (C) and (D) are defined using the Harvard-Oxford cortical structural atlas with a probabilistic threshold of 30%. All four masks were used for small volume correction (SVC) in the univariate analysis. Masks (A) and (B) were also used in the ROI-based multivariate RSA analysis.

